# Variations in ERP Data Quality Across Paradigms, Participants, and Scoring Procedures

**DOI:** 10.1101/2022.08.22.504647

**Authors:** Guanghui Zhang, Steven J. Luck

**Affiliations:** Center for Mind and Brain, University of California–Davis, Davis, California, 95618, USA

**Keywords:** ERP, data quality, standardized measurement error, number of trials, trial-to-trial EEG variability, scoring methods

## Abstract

Although it is widely accepted that data quality for event-related potential (ERP) components varies considerably across studies and across participants within a study, ERP data quality has not received much systematic analysis. The present study used a recently developed metric of ERP data quality— the standardized measurement error (SME)—to examine how data quality varies across different ERP paradigms, across individual participants, and across different procedures for quantifying amplitude and latency values. The EEG recordings were taken from the ERP CORE, which includes data from 40 neurotypical college students for seven widely studied ERP components: P3b, N170, mismatch negativity, N400, error-related negativity, N2pc, and lateralized readiness potential. Large differences in data quality were observed across the different ERP components, and very large differences in data quality were observed across participants. Data quality also varied depending on the algorithm used to quantify the amplitude and especially the latency of a given ERP component. These results provide an initial set of benchmark values that can be used for comparison with previous and future ERP studies. They also provide useful information for predicting effect sizes and statistical power in future studies, even with different numbers of trials. More broadly, this study provides a general approach that could be used to determine which specific experimental designs, data collection procedures, and data processing algorithms lead to the best data quality.

## 1. Introduction

Event-related potentials (ERPs) are typically quite small relative to the background noise. For example, the face-sensitive N170 component might have an amplitude of 4 μV but might be embedded in 40 μV of background EEG. Conventionally, researchers average multiple trials together to isolate the ERP and “average out” the noise. The amplitude and/or latency of a given component is then quantified or scored from the averaged ERP waveforms. Finally, these scores are entered into a statistical analysis to compare experimental conditions or groups of participants. Other approaches are also common in ERP research, but this averaging-followed-by-scoring sequence is the dominant approach in many subfields ^1^.

Because ERPs are so small relative to the background noise, the averaged ERP waveforms often contain considerable noise that adds uncontrolled variability to the observed amplitude and latency scores. This uncontrolled variability carries forward to increase the variance across participants, reducing effect sizes and the statistical power for detecting differences among conditions or groups. Although it is widely appreciated that noisy ERPs are problematic, and that averaged ERP waveforms are much noisier in some paradigms and participants than in others, there is no widely used metric of data quality in ERP research for quantifying this noise ^2^.

### 1.1. The standardized measurement error as a metric of ERP data quality

Recently, we proposed a metric of data quality for averaged ERPs called *standardized measurement error* (SME) (Luck et al., 2021). The SME is a special case of the standard error of measurements, and it is designed to quantify the precision of measurements (e.g., the amplitude or latency scores) that are obtained from averaged ERP waveforms. As detailed by Brandmaier et al. (2018), a measure is precise to the extent that the same value is obtained upon repeated measurements^3^, assuming that the measurement process does not influence the system being measured. In theory, the precision of an ERP amplitude or latency score for a given participant could be quantified by repeating the experiment a large number of times with that participant, obtaining the score for each repetition of the experiment, and computing the standard deviation (SD) of these scores. However, this would be unrealistically time-consuming in practice, and the ERPs would likely change over repetitions of the experiment as a result of learning, boredom, etc.

Fortunately, it is possible to estimate the precision of an ERP score using the data from a single recording session. This is particularly straightforward when the score being obtained is the mean amplitude over some time range (e.g., the mean voltage from 350-550 ms for the P3b component, as illustrated in Figure 1). The mean amplitude score obtained from an averaged ERP waveform is identical to the average of the mean amplitude scores obtained from the single-trial EEG epochs, so the standard error of measurement for a mean amplitude score is simply the standard error of the mean or SEM of the score. A widely-used analytic solution is available for estimating the SEM:

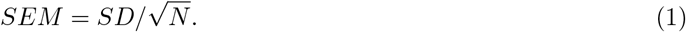

**Figure 1.**
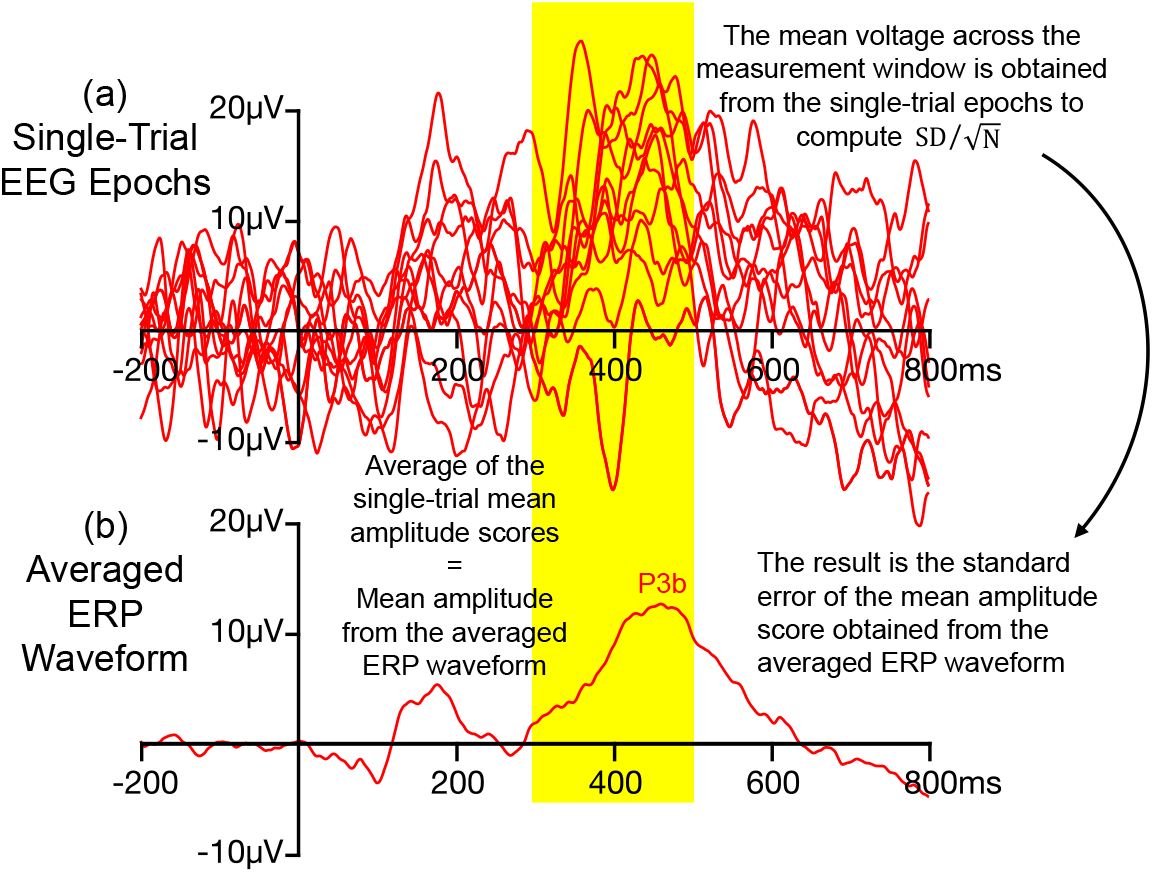
Example of how Equation 1 is used to estimate the standard error of measurement when the amplitude of the P3b wave is scored from an averaged ERP waveform as the mean voltage across a measurement window of 350-550 ms. To compute this standard error, the mean amplitude score is obtained from the single-trial EEG epochs, and these single-trial scores are used to compute 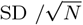. SD is the standard deviation of the single-trial mean amplitude scores, and N is the number of trials. The result is the standard error of measurement for the score obtained from the averaged ERP. Separate standard error values can be obtained for each experimental condition (e.g., rare trials versus frequent trials in an oddball paradigm). This approach is possible because the mean amplitude score obtained from the averaged ERP waveform is equal to the average of the single-trial mean amplitude scores. This approach does not work with other scoring methods, such as peak amplitude and peak latency. Note that only a subset of the single-trial epochs used to create the averaged ERP waveform are shown here.

In this equation, is the standard deviation of the single-trial mean amplitude scores and N is the number of trials being averaged together to create the averaged ERP waveform. This is illustrated in Figure 1, which shows single-trial EEG epochs and the corresponding averaged ERP waveform from a hypothetical oddball experiment in which the P3b component is scored as the mean voltage between 350 and 550 ms. The standard error of this score is estimated by measuring the mean voltage from 350-550 ms in the single-trial EEG epochs, taking the SD of these values, and dividing by the square root of N. In other words, although the ultimate amplitude score is obtained from the averaged ERP waveform, the standard error of this score is obtained by applying Equation 1 to measurements obtained from the single-trial EEG epochs. The resulting standard error is an estimate the precision of the mean amplitude score that is obtained from the averaged ERP waveform (see Luck et al. (2021) for a more detailed explanation and justification).

When other scoring methods are used, the score obtained from the averaged ERP waveform is not equal to the average of the single-trial scores, so Equation 1 cannot be used to estimate the standard error of measurement for these scores. For example, if you obtain the peak amplitude from the single-trial epochs in Figure 1a and then average these values together, the result will not be equal to the peak amplitude measured from the averaged ERP waveform in Figure 1b. Thus, Equation 1 cannot be used to estimate the standard error of the peak amplitude score obtained from an averaged ERP waveform. However, bootstrapping can be used to estimate the standard error of measurement for the peak amplitude or for virtually any other amplitude or latency score that is obtained from averaged ERP waveforms. In this approach, the set of individual trials that were actually collected for a given participant are used to provide a population of trials for simulating repetitions of the experiment for that participant. A single recording session is then simulated by randomly sampling from this set of trials. By simulating a large number of sessions, and measuring the amplitude or latency score of interest from each of these simulations, it is possible to quantify the variability of scores across simulated sessions and obtain an estimate of the standard error of measurement (for more details, see section 2.4.2 and Luck et al. (2021)). Another advantage of bootstrapping is that it can be used when the score is obtained from a transformation of the averaged ERP waveforms, such as a difference wave.

Whereas Equation 1 involves dividing by the square root of the number of trials 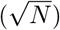, bootstrapping does not explicitly involve this division step. Nonetheless, because bootstrapping and Equation 1 are just two different ways of estimating the same value, standard errors vary with the number of trials in approximately the same way whether they are estimated using bootstrapping or using Equation 1.

When the standard error of measurement is used to quantify the precision of an ERP amplitude or latency score, we refer to it as the standardized measurement error (SME) of that score. It estimates how much variability would be present in the mean amplitude scores from a given participant if the experiment were repeated an infinite number of times (assuming no learning, fatigue, etc.) and the amplitude or latency score was obtained from the averaged ERP waveform on each repetition. Specifically, the SME is an estimate of the SD of the scores we would get across these hypothetical repetitions. When Equation 1 is used to estimate the SME for mean amplitude scores, we call this the analytic SME (aSME); when bootstrapping is used, we call this the bootstrapped SME (bSME).

As detailed by Luck et al. (2021), the SME can be used to determine how much of the variability across participants in amplitude or latency scores is a result of measurement error versus true differences among participants. This makes it possible to determine the extent to which the effect size for a comparison between groups or conditions is impacted by measurement error. It can also be used to predict exactly how effect sizes and statistical power will change if the measurement error is increased or decreased (e.g., by changing the number of trials per participant). The SME can be computed using the ERPLAB Toolbox software package (Lopez-Calderon & Luck, 2014), beginning with Version 8.

### 1.2. Defining “data quality”

The SME is intended to be a metric of data quality. Luck et al. (2021) argued that the concept of data quality in ERP research must be defined with respect to the specific scores that will be obtained from the averaged ERP waveforms. This is because the impact of a given source of noise will depend on how the amplitude or latency of a component is being scored. For example, high-frequency noise has a large effect on peak amplitude scores but relatively little effect on mean amplitude scores (because the upward and downward deflections produced by high-frequency noise largely cancel out when the voltages are averaged over a broad measurement window). Similarly, low-frequency drifts have a large impact on data quality for the amplitude of late ERP components such as the P3b or N400 (Kappenman & Luck, 2010; Tanner et al., 2015), but these drifts have less impact for earlier components (because the signal has not had much time to drift betewen the baseline period and the measurement period). Thus, there is no meaningful definition of ERP data quality that is independent of the scoring method.

When the SME is used to quantify data quality, any factors that produce uncontrolled variability in a given amplitude or latency score are considered to be noise with respect to that specific score. This definition of noise includes factors that might be of considerable theoretical or practical interest, such as oscillations that are not phase-locked to stimulus onset (Busch et al., 2009; Mathewson et al., 2009) or trial-to-trial variations in attentional state (Adrian & Matthews, 1934; Boudewyn et al., 2017). Indeed, a great deal of evidence indicates that trial-to-trial variations in neurocognitive processes are important for understanding both typical and atypical cognitive processing (Ratcliff & McKoon, 2008; Tamm et al., 2012). However, these factors are considered noise from the perspective of the data quality for a given amplitude or latency score because they decrease the precision of the score and therefore decrease effect sizes and statistical power. In addition, because SME values combine sources of variability that are functionally important (e.g., variations in attentional state) and sources of variability that play no functional role (e.g., induced electrical noise), the SME is not appropriate for use as a measure of trial-to-trial variability in neurocognitive processing.

It is also important to note that the SME is directly influenced by the number of trials (N), and differences in N across studies, conditions, or participants will lead to differences in SME. This is appropriate, because the SME is designed to quantify the precision of the amplitude or latency scores that will be entered into the statistical analysis, and differences in N will impact the precision of these scores and the resulting statistical power. However, it may sometimes be useful to compare data quality values in a manner that is not dependent on the number of trials and instead purely reflects trial-to-trial variability in the EEG. This is trivial to accomplish when Equation 1 is used to compute aSME values for mean amplitude scores, because the SD of the single-trial scores can be used to estimate the trial-to-trial variability. We therefore provide these SD values along with SME values in the main analyses.

Unfortunately, this approach is not possible for bootstrapped SME values, because bootstrapping does not directly provide a measure of the trial-to-trial variability. We are currently developing a solution for this (Zhang & Luck, in preparation).

### 1.3. Potential uses of the SME

The SME has many potential uses. Within a given study, the SME could be used to determine which participants are so noisy that they should be excluded, which channels are so noisy that they should be interpolated, and how changing a given processing parameter (e.g., the artifact rejection threshold) will increase or decrease the data quality. When new laboratories are built or new personnel are trained, the SME makes it possible to determine whether the resulting data quality meets an objective standard. In methodology research, the SME could be used to determine which recording and analysis procedures lead to the highest data quality. If published papers regularly reported SME values, it would be possible to quantitively assess how data quality varies among different experimental paradigms, subject populations, and processing pipelines.

For many of these uses, it would be valuable to have a broad set of benchmark SME values against which new data could be compared. That is, it would be useful to have a reference point that can be used to make an informed guess about the range of SME values that should be expected in a given study. The primary goal of the present paper was therefore to provide an initial set of benchmark values. Specifically, we computed SME values for four different scoring procedures (peak amplitude, mean amplitude, peak latency, and 50% area latency) obtained from each of 40 participants for each of the seven ERP components contained in the ERP CORE (Compendium of Open Resources and Experiments; Kappenman et al. (2021)). The ERP CORE is a set of stimulus presentation scripts, data analysis scripts, and EEG recordings for six standard ERP paradigms that yield seven commonly studied ERP components. Each task requires approximately 10 minutes to run, and the resource contains data from 40 neurotypical college students who completed all six tasks in a single session.

Because the ERP CORE contains data from a broad range of paradigms and a reasonably large set of participants, it provided an excellent resource for developing an initial set of benchmark SME values. Well-controlled comparisons across ERP components were possible because the same participants completed all six paradigms, and a broad variety of experimental details were held constant across paradigms (e.g., lighting, viewing distance). Moreover, this resource made it possible to determine the extent to which data quality is correlated across components and scoring methods (e.g., whether an individual with poor data quality for one component or scoring method also has poor data quality for other components or scoring methods).

Of course, data quality is likely to differ between the neurotypical college students who were tested for the ERP CORE and other populations (e.g., children, adults with neurological or psychiatric disorders). Thus, the SME values from the ERP CORE data provide a direct benchmark only for populations that are similar to the participants that are included in the ERP CORE. For other populations, the ERP CORE data are not a benchmark per se, but they are still useful for providing a point of comparison until benchmark values can be obtained for those populations.

### 1.4. Organization of the present paper

The SME analyses in the present paper are divided into three sections. The first section provides a detailed description of the SME values obtained from the ERP CORE data across paradigms, across participants, and across scoring procedures. These values are presented in multiple different formats to make it easy to compare them with SME values obtained in future studies. The second section asks why the SME values varied across the paradigms and participants, focusing on the number of trials being averaged together and trial-to-trial variability in the EEG. The final section of the present paper quantifies the extent to which SME values for a given individual are correlated across paradigms and across scoring procedures. In other words, this section asks whether a given individual has generally “good” or “bad” data quality across paradigms and scoring procedures. Together, these analyses provide a useful starting point for researchers who wish to examine the data quality in their own paradigms, participants, and scoring procedures.

## 2. Method

All the scripts and results for the present analyses have been added to a folder named SME in the online repository for the ERP CORE (https://doi.org/10.18115/D5JW4R). This includes spreadsheets with all the single-subject SME values. The participants, experimental paradigms, recording methods, and analysis methods are described in detail in Kappenman et al. (2021). Here, we provide a brief overview.

### 2.1. Participants

Data were obtained from 40 neurotypical college students (25 female) from University of California, Davis community. Although some participants failed to meet the inclusion criteria for some of the components in the original ERP CORE analysis (e.g., owing to poor behavioral performance), we provide SME data from all 40 participants here to represent the entire range of data quality across participants. The one exception was that Subject 7 was excluded from the N2pc analyses because the number of trials for one condition was zero, making it impossible to compute SME values for this participant in this paradigm.

### 2.2. Overview of the six paradigms and seven ERP components

Figure 2 provides an overview of the six paradigms, and Figure 3 shows the grand average parent waves (left panel) and difference waves (right panel) for the seven ERP components. These grand average waveforms are identical to those provided by Kappenman et al. (2021), except that the waveforms here include all participants (except for Subject 7 in the N2pc paradigm, who had zero trials in one condition). By contrast, Kappenman et al. (2021) excluded participants who exceeded criteria for the percentage of trials rejected because of artifacts or behavioral errors. Because the goal of the present study was to assess the entire range of data quality, which is impacted by the number of rejected trials, the present analyses included all participants (except Subject 7, for whom data quality was undefined in the N2pc experiment).

**Figure 2.**
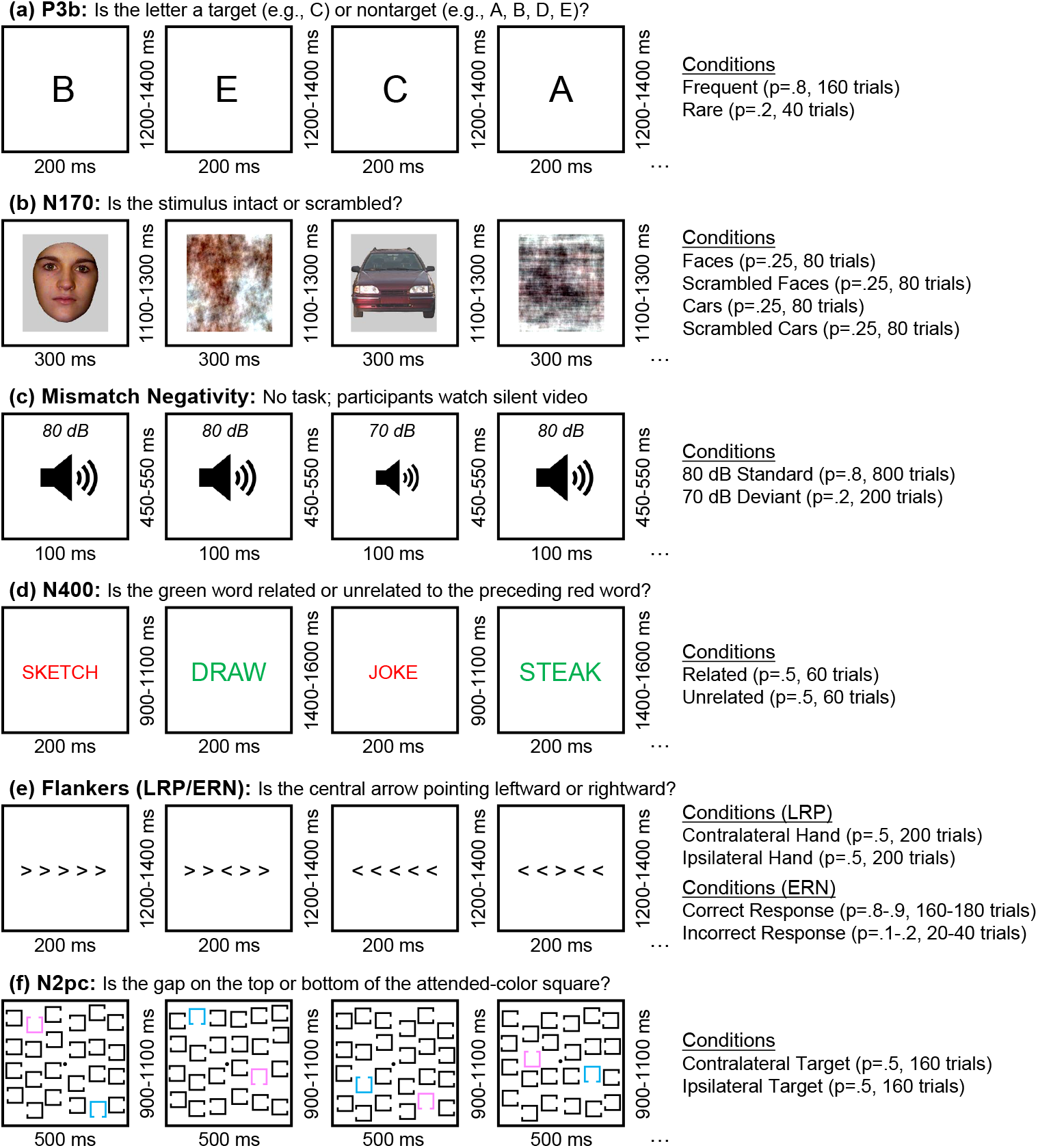
Examples of multiple trials for each of the six paradigms. (a) Active visual oddball paradigm used to elicit the P3b component. The letters A, B, C, D, and E were presented in random order (p = .2 for each letter). In each block, one letter was designated the target, and the other four letters were nontargets. Participants were required to classify each stimulus as target (20% of stimuli) or non-target (80% of stimuli). (b) Face perception paradigm used to elicit the N170 component. On each trial, a stimulus from one of four equiprobable categories was displayed (face, scrambled face, car, scrambled car), and participants were required to classify the image as an intact object (face or car) or a texture (scrambled face or scrambled car). The present paper focuses only on the face and car trials. (c) Passive auditory oddball task used to elicit the mismatch negativity (MMN). On each trial, either a standard tone (80 dB, p = .8) or a deviant tone (70 dB, p = .2) was presented. The tones were task-irrelevant; participants watched a silent video during this paradigm. (d) Word pair judgment paradigm used to elicit the N400 component. On each trial, a red prime word was followed by a green target word, and participants indicated whether the green word was related (p = .5) or unrelated (p = .5) to the preceding red prime word. (e) Flankers task used to elicit the lateralized readiness potential (LRP) and the error-related negativity (ERN). Participants were required to indicate whether the central arrow pointed leftward or rightward, ignoring the flanking arrows. (f) Simple visual search task used to elicit the N2pc component. One color (pink or blue) was designated the target color at the beginning of each trial block. On each trial, participants indicated whether the gap was on the top or the bottom of the attended-color square.

**Figure 3.**
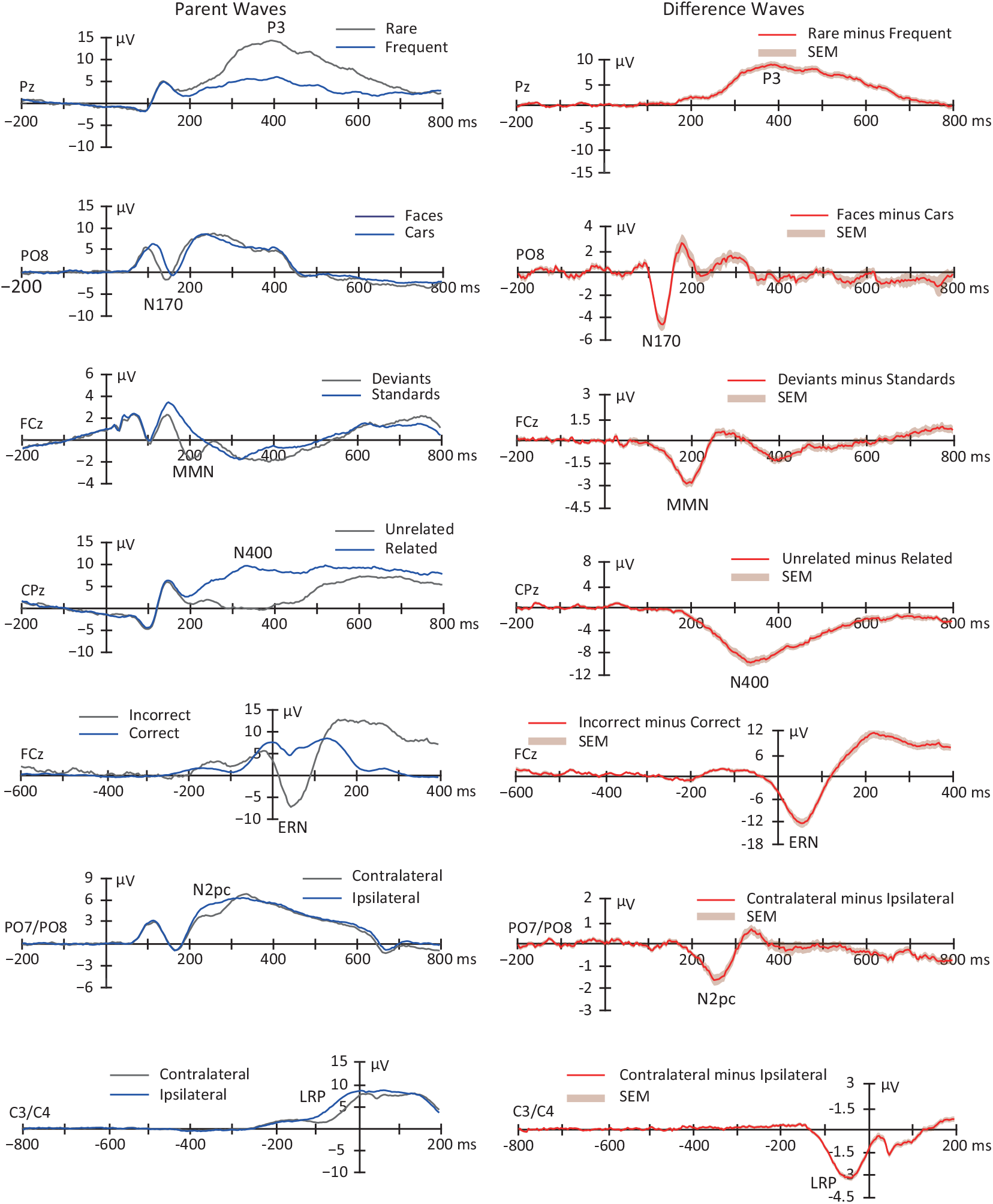
Grand average parent waves (left panel) and difference waves (right panel) for the seven ERP components examined in the ERP CORE]. The shaded area accompanying the difference waves is the standard error of the mean across participants at each time point. This is largely identical to Figure 2 in Kappenman et al. (2021), except that all 40 participants were included here (except that Subject 7 was excluded from the N2pc waveforms because the number of trials for one condition was zero after artifact rejection). The original figure was released under a CC BY license (http://creativecommons.org/licenses/by/4.0/).

The P3b paradigm is shown in Figure 2 a. In each block of this visual oddball task, five letters (A, B, C, D, and E) appeared in random order (p = .2 for each letter). One letter was designed to be target for a given block and the other four letters were non-targets (e.g., “A” was the target in one block and a nontarget in the other blocks). Participants were instructed to press one button if a given letter was the target and a different button if it was one of the four nontargets.

The N170 paradigm is shown in Figure 2 b. Each trial consisted of a face, a car, a scrambled face, or a scrambled car (p = .25 for each category). For each stimulus, participants pressed one of two buttons to indicate whether the stimulus was an “intact object” (regardless of whether it was a face or car) or a “texture” (scrambled face or scrambled car). For the sake of simplicity, we examined only the face and car trials. Face and car stimuli were modified from (Rossion & Caharel, 2011).

The mismatch negativity (MMN) paradigm is shown in Figure 2 c. In this passive auditory oddball paradigm, a task-irrelevant sequence of standard tones (80 dB, p = .8) and deviant tones (70 dB, p = .2) was presented to participants while they watched a silent video. No responses were made to the tones.

The N400 paradigm is shown in Figure 2 d. In this word pair judgment task, each trial consisted of a red prime word followed by a green target word. On each trial, participants were required to press one of two buttons to indicate whether the target word was related (p = .5) or unrelated (p = .5) to the preceding prime word.

The paradigm used to examine the lateralized readiness potential (LRP) and the error-related negativity (ERN) is shown in Figure 2 e. In this variant of the Eriksen flanker paradigm (Eriksen & Eriksen, 1974), each stimulus contained a central arrow surrounded by flanking arrows that pointed in the same direction or the opposite direction as the central arrow. One each trial, participants pressed one of two buttons to indicate whether the central arrow was pointing leftward (p = .5) or rightward (p = .5).

The N2pc paradigm is shown in Figure 2 f. In this simple visual search task, either pink or blue was designed the target color for a given block of trials. Each stimulus within a block contained a pink square, and blue square, and 22 black squares. For each stimulus, participants pressed one of two buttons to indicate the location (top or bottom) of a gap in the attended-color square.

### 2.3. Overview of data collection and data processing pipeline

Continuous EEG data were collected using a Biosemi ActiveTwo recoding system with active electrodes (Biosemi B.V., Amsterdam, the Netherlands) an antialiasing filter (fifth order sinc filter with a half-power cutoff at 204.8 Hz) and a sampling rate of 1024 Hz. Data were analyzed from 30 scalp sites along with horizontal and vertical electrooculogram electrodes.

The present analyses were performed on the preprocessed data provided as part of the ERP CORE resource (https://doi.org/10.18115/D5JW4R). Our goal was to examine data quality in the context of the kind of preprocessing that would typically be performed in an ERP study (e.g., filtering, referencing, artifact rejection and correction), so we used the data files that were already preprocessed (see Luck (2022) for examples of how preprocessing influences the SME). The original preprocessing and our additional analyses were conducted in MATLAB 2020a environment using the EEGLAB 2021.1 (Delorme & Makeig, 2004) and ERPLAB 8.30 (Lopez-Calderon & Luck, 2014) . All scripts are available in the ERP CORE resource.

The preprocessing steps are described in detail by Kappenman et al. (2021), and here we provide a brief summary. The event codes were shifted to reflect the intrinsic delay of the video monitor, and the data were resampled at 256 Hz. The data were referenced to the average of the P9 and P10 electrodes (close to the left and right mastoids) for all components except the N170, for which the average of all scalp sites was used as the reference. A noncausal Butterworth high-pass filter (half-amplitude cutoff 0.1 Hz, 12dB/oct roll-off) was applied. Independent component analysis (ICA) was used to correct the data for eyeblinks and eye movements.

The resulting EEG data were epoched and then baseline-corrected using the time windows shown in Table 1. Bad channels were interpolated using ERPLAB’s spherical interpolation algorithm. Trials with blinks or eye movements that could have impacted perception of the stimuli were rejected, as were trials with large EEG deflections in any channel and trials with incorrect behavioral responses. The remaining epochs were averaged across trials for each experimental condition.

**Table 1.** Epoch window, baseline period, electrode site, and time window used for each ERP component.

| Component | P3b | N170 | MMN | N400 | ERN | N2pc | LRP |
| --- | --- | --- | --- | --- | --- | --- | --- |
| Epoch window (ms) | -200 to 800 | -200 to 800 | -200 to 800 | -200 to 800 | -600 to 400 | -200 to 800 | -800 to 200 |
| Baseline period (ms) | -200 to 0 | -200 to 0 | -200 to 0 | -200 to 0 | -400 to -200 | -200 to 0 | -800 to -600 |
| Electrode site | Pz | PO8 | FCz | CPz | FCz | PO7/PO8 | C3/C4 |
| Time window (ms) | 300 to 600 | 110 to 150 | 125 to 225 | 300 to 500 | 0 to 100 | 200 to 275 | -100 to 0 |

Table 2 shows the mean number of epochs remaining for averaging in each condition for each ERP component, and the ERP CORE repository (https://doi.org/10.18115/D5JW4R) includes spreadsheets with the number of included and excluded trials for each participant. Table 2 also includes the mean and peak amplitudes for each ERP component.

**Table 2.** Mean number of trials (±SEM) and mean of mean/peak amplitude (±SEM) across all 40 participants for each condition for the seven ERP components, after excluding trials with artifacts and behavioral errors.

|  | P3b |  | N170 |  | MMN |  | N400 |  |
| --- | --- | --- | --- | --- | --- | --- | --- | --- |
|  | Rare | Frequent | Faces | Cars | Deviants | Standards | Unrelated | Related |
| #Trials | 30.53 ± 1.36 | 139.90±3.29 | 69.00 ±2.04 | 68.13 ±1.44 | 183.18 ± 3.39 | 534.95± 9.51 | 52.60 ±1.08 | 51.20 ±1.18 |
| Mean amplitude | 11.39 ± 0.79 | 4.34±0.46 | 0.54 ±0.85 | 4.16 ±0.79 | 0.13 ± 0.23 | 1.99± 0.19 | 1.01 ±0.61 | 8.72 ±0.83 |
| Peak amplitude | 17.44 ± 1.03 | 8.62±0.58 | -2.98 ±0.88 | 0.94 ±0.94 | -2.50 ± 0.29 | 0.56± 0.34 | -2.89 ±0.63 | 4.88 ±0.85 |
|  | ERN |  | N2pc |  | LRP |  | – |  |
|  | Incorrect | Correct | Left target | Right target | Left response | Right response | – | – |
| #Trials | 40.05 ±3.42 | 337.03± 7.39 | 118.73 ±4.33 | 118.23± 4.61 | 166.75 ±3.94 | 167.13 ± 3.78 | – | – |
| Mean amplitude | -2.97 ±1.00 | 6.27± 0.84 | 3.62 ±0.52 | 4.79± 0.53 | 3.27 ±0.65 | 5.75 ± 0.64 | – | – |
| Peak amplitude | -9.17 ±1.20 | 3.87± 0.92 | 2.62 ±0.60 | 3.42± 0.56 | 0.49 ±0.65 | 3.04 ± 0.74 | – | – |

### 2.4. Quantification of data quality

#### 2.4.1. Measurement windows and electrode sites

The original ERP CORE paper (Kappenman et al., 2021) identified an optimal electrode site and an optimal time window for scoring each ERP component, which are shown in Table 1. We used these sites and windows for obtaining the amplitude and latency scores and for quantifying the SME and trial-to-trial variability of these scores.

#### 2.4.2. SME quantification

We focused on four scoring algorithms, as implemented by ERPLAB Toolbox (Lopez-Calderon & Luck, 2014). Mean amplitude was scored as the mean voltage across time points within the measurement window for a given component. Peak amplitude was scored as the voltage of the most positive point (for P3b) or most negative point (for the other components) within the measurement window. Peak latency was scored as the latency of peak amplitude point ^4^. The 50% area latency was scored by measuring the area of the region above the zero line (for P3b) or below the zero line (for the other components) within the measurement window, and then finding the time point that bisected this area into two equal-area regions. To increase precision, the waveforms were upsampled by a factor of 10 using spline interpolation before the latencies were scored (see Luck (2014) for the rationale).

According to Equation 1, the analytic SME (aSME) for mean amplitude can be estimated by measuring the mean amplitude on single trials and dividing the standard deviation of the single-trial amplitudes by the square root of the number of trials. However, this approach is not valid for peak amplitude, peak latency and 50% area latency, and it cannot be directly applied to difference waves. We therefore computed the bootstrapped SME (bSME), even for mean amplitude scores. Note that aSME and bSME values are virtually identical for mean amplitude scores as long as the number of trials is reasonably large (more than eight).

These bSME values were obtained for each of the parent waves used to define a given component (e.g., the rare and frequent trials in the P3b paradigm) and also for the corresponding difference wave (e.g., the rare-minus-frequent difference wave in the P3b paradigm). As will be described in Section 3.2, the latency scores could not be validly obtained from the parent waveforms in many cases, so latencies were obtained only from the difference waves.

Bootstrapping is a common approach in many areas of statistics (Boos, 2003; Efron & Tibshirani, 1994). As described by Luck et al. (2021), we implemented bootstrapping by simulating 1000 repetitions of each experiment for each participant. In each simulated repetition of a given experiment, we selected N trials at random, with replacement, from all N trials that were used to create the standard averaged ERP waveforms for a given condition in that participant. Remarkably, sampling with replacement from the existing set of N trials accurately simulates conducting a replication experiment with N new trials as long as N is reasonably large (e.g., 8; Chernick (2011)). We then averaged that set of N trials together and obtained the mean and peak amplitude scores from the averaged ERP waveform. The SME was calculated as the SD across the 1000 simulated repetitions for that condition in that participant.

For each repetition, we also created a difference wave for the two conditions of a given experiment. We then obtained the mean amplitude, peak amplitude, peak latency, and 50% area latency scores from this difference wave. The SME for a given difference-wave score was then computed as the SD of the scores from the 1000 simulated repetitions of the experiment.

One limitation of this bootstrapping procedure is that, because it involves sampling randomly from the available trials, the SME value varies slightly each time the procedure is repeated. To make the results exactly reproducible (e.g., if another lab wishes to reproduce the results), a random number generator seed can be generated for each iteration and then used across repetitions of the procedure.

#### 2.4.3. Quantification of trial-to-trial variability

For mean amplitude scores, Equation 1 indicates that trial-to-trial variability—quantified as the SD of the single-trial mean amplitudes—is a key factor in determining measurement error. We therefore computed the SD of the single-trial mean amplitudes for the parent waves in each experimental condition for each ERP component for each participant. This was straightforward for the P3b, N170, MMN, N400, and ERN components, but it was slightly more complicated for the N2pc and LRP because the parent waves were defined as contralateral (the left hemisphere signal for trials with a right-side stimulus or response averaged with the right hemisphere signal for trials with a left-side stimulus or response) or ipsilateral (the left hemisphere signal for trials with a left-side stimulus or response averaged with the right hemisphere signal for trials with a right-side stimulus or response). To obtain the SD for the contralateral and ipsilateral parent waveform, we took advantage of the fact that the variance of a sum of two random variables is equal to the sum of the two variances. Specifically, we computed the variance for each of the two waveforms that were combined (e.g., the variance of the left hemisphere mean amplitudes for trials with a left-side stimulus or response and the variance of the right hemisphere mean amplitudes for trials with a right-side stimulus or response), took the average of these two variances, and then took the square root to yield an SD value.

We also computed SD values for the single-trial peak amplitude scores. Single-trial latency scores could not be validly computed for several components, so we did not examine trial-to-trial variability in latency. That issue will be addressed via simulations in a subsequent paper (Zhang & Luck, in preparation).

#### 2.4.4. Quantification of trial-to-trial variability

We used F and t tests to compare SME and SD values across scoring methods and experimental paradigms. We used Spearman rho rank-order correlation coefficients to examine how SME or SD values covaried across participants for different scoring methods or experimental paradigms. Slope values, however, were obtained from standard linear regressions. For each set of statistical analyses, we performed a familywise correction for multiple comparisons using the false discovery rate correction (Benjamini & Hochberg, 1995). An alpha of .05 was used in all statistical tests.

## 3. Results

### 3.1. Basic characterization of data quality across paradigms, participants, and scoring procedures

We begin by providing basic information about how SME values varied across the seven ERP components, the two main conditions used to isolate each component, the four different amplitude and latency scoring procedures, and the 40 different participants. A large number of SME values are presented. To keep things manageable, the key values are summarized in the tables and figures of the main manuscript, and additional values are provided in supplementary tables and figures. In addition, spreadsheets containing the single-participant values are available online at https://doi.org/10.18115/D5JW4R, along with all the codes used to compute the SME values.

#### 3.1.1. Variations in data quality across paradigms, conditions, and scoring procedures (parent waves)

Figure 4 a shows the SME for the mean amplitude scores, averaged across participants, for each of the parent waves used to define the seven components. Figure 4 b shows the corresponding SME values for the peak amplitude scores. The exact values are presented in supplementary Table S1. The root mean square (RMS) of the SME values across participants is sometimes more useful than mean SME across participants ^5^, and the RMS values are provided in supplementary Figure S1 and supplementary Table S2. Figures 4 c-f show the variability (SD) across trials and the square root of these number of trials; these will be discussed in Section 3.2. Note that it is difficult to obtain valid latency measures from parent waveforms in many cases. For example, there is no negative-going voltage deflection on semantically related trials in the N400 paradigm (see Figure 3), so N400 latency cannot be validly measured for this experimental condition. Thus, this subsection focuses on the SME for the amplitude scores obtained from parent waveforms.

**Figure 4.**
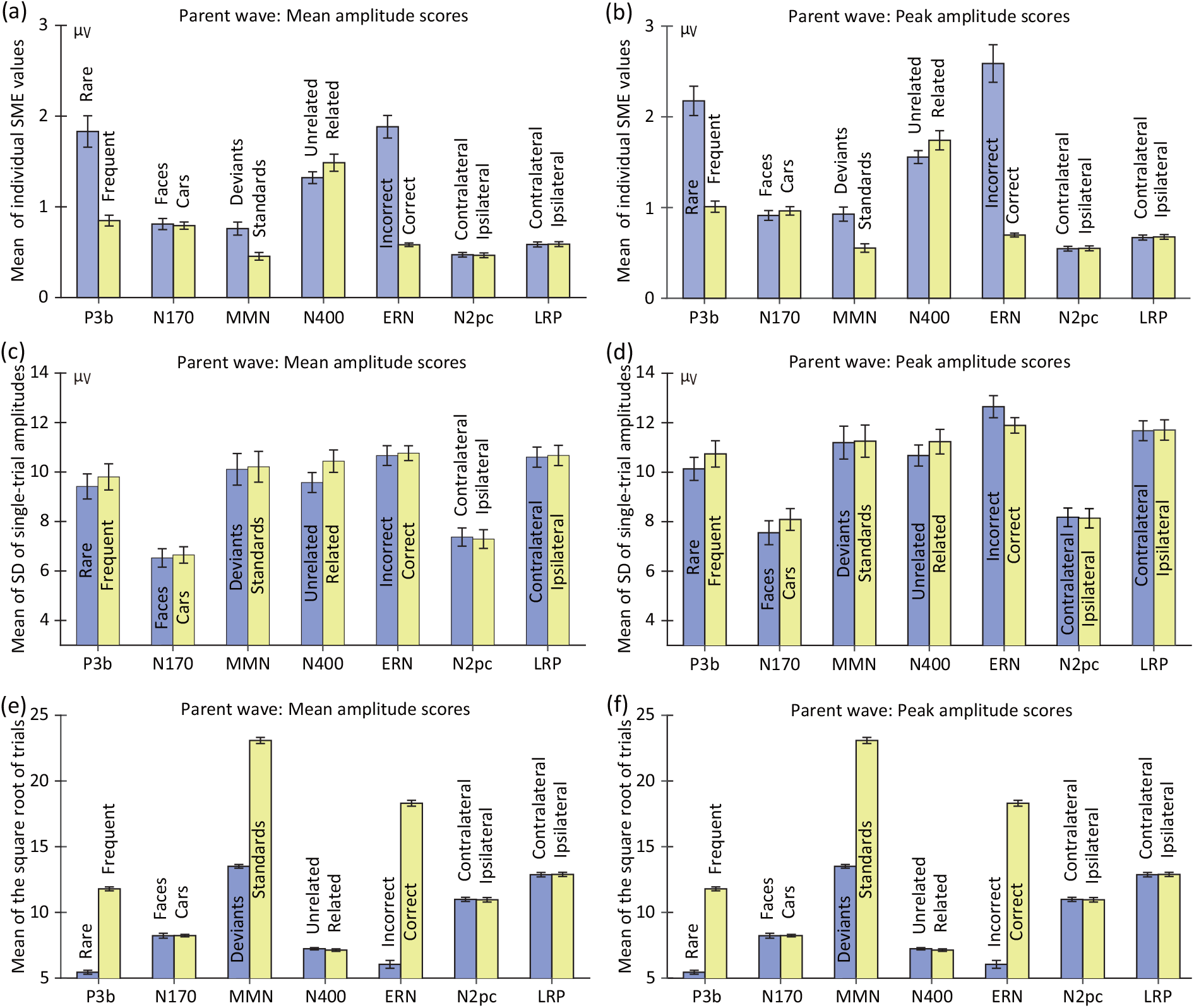
Mean across participants of the standardized measurement error (SME; panels a and b), the standard deviation across trials (SD; panels c and d), and the square root of the number of trials (panels e and f). Separate values are provided for the SME values corresponding to mean amplitude scores (left column) and peak amplitude scores (right column). The mean and peak amplitude scores and corresponding SME and SD values were computed using the time windows and electrode sites shown in Table 1. Note that the number of trials varied across individuals because of artifact rejection and exclusion of trials with errors. However, panels e and f are identical because the same trials were used for the averages used for scoring mean amplitude and peak amplitude. Error bars show the standard error of the mean of the single-participant values.

The first thing to note in Figure 4 is that the mean SME values were worse (higher) for the peak amplitude scores than for the mean amplitude scores in every case. To test this statistically, we used paired t tests to compare the SME values for mean amplitude and peak amplitude, separately for each combination of experimental paradigm and condition (correcting for multiple comparisons). In all 14 cases, the SME was significantly worse (higher) for peak amplitude than mean amplitude (see Table S3). This finding is consistent with the claim that peak amplitude is more sensitive to noise than mean amplitude (Clayson et al., 2013; Luck, 2014).

The next thing to note is that the SME values varied considerably across the seven ERP components, with some components having much worse (higher) SME values than other components. For example, the SME for mean amplitude (measured from the difference waves) was approximately four times greater for P3b and N400 than for N2pc. The variations in SME across components were even more extreme for the peak amplitude scores. In addition, the SME values varied considerably between the two conditions used to define some of the components (e.g., much higher for the rare category than for the frequent category in the P3b paradigm). The reasons for these differences will be examined in Sections 3.2 and 3.3.

To analyze these differences statistically, we used paired t tests to compare each pair of conditions (correcting for multiple comparisons). As shown in Table 3, conditions with fewer trials yielded significantly higher SME values than conditions with more trials (i.e., for the P3b, MMN, and ERN scores). In addition, the SME in the N400 paradigm was significantly greater for the semantically related condition than for the semantically unrelated condition. A possible explanation for this difference will be described in Section 4.2.

**Table 3.** Paired t tests comparing either the standardized measurement error (SME) or the standard deviation across trials (SD) between the two experimental conditions for each ERP component, separately for mean amplitude and peak amplitude (corrected for multiple comparisons across the family of tests for each scoring method). Participant 7 was excluded from the N2pc analyses (but not for the other analyses) because the number of trials was zero in one of the N2pc conditions for this participant.

| Parent wave | P3b |  | N170 |  | MMN |  | N400 |  | ERN |  | N2pc |  | LRP |  |
| --- | --- | --- | --- | --- | --- | --- | --- | --- | --- | --- | --- | --- | --- | --- |
|  | t(39) | p | t(39) | p | t(39) | p | t(39) | p | t(39) | p | t(38) | p | t(39) | p |
| SME for mean amplitude | 7.79 | <.001 | 0.36 | 0.723 | 10.19 | <.001 | -3.29 | 0.004 | 10.75 | <.001 | 0.99 | 0.459 | -0.71 | 0.560 |
| SME for peak amplitude | 10.23 | <.001 | -1.26 | 0.302 | 11.40 | <.001 | -3.41 | 0.004 | 9.21 | <.001 | -0.51 | 0.614 | -0.95 | 0.405 |
| SD for mean amplitude | -1.76 | 0.301 | -0.67 | 0.594 | -1.17 | 0.487 | -3.43 | 0.007 | -0.41 | 0.686 | 0.95 | 0.487 | -1.07 | 0.487 |
| SD for peak amplitude | -2.27 | 0.102 | -1.94 | 0.103 | -0.60 | 0.706 | -1.97 | 0.103 | 2.34 | 0.102 | 0.38 | 0.706 | -1.56 | 0.706 |

Because there were two conditions in each paradigm, there was no straightforward way to statistically compare SME values across paradigms in a condition-independent manner for the SME values shown in Figure 4. However, the next section provides a comparison across paradigms for amplitude and latency scores obtained from difference waves.

#### 3.1.2. Variations in data quality across paradigms and scoring procedures for experimental effects (difference waves)

In many cases, it is useful to score the amplitude or latency of a component from a difference wave (Luck, 2014). The SME for such scores can be obtained using bootstrapping (Luck et al., 2021). For mean amplitude scores, the resulting SME quantifies the measurement error of the experimental effect. Difference waves are also necessary for obtaining valid latency scores for some components (e.g., N2pc and LRP). Thus, this subsection focuses on scores obtained from difference waves, which made it possible to characterize the SME for both amplitude measures (mean amplitude and peak amplitude) and latency measures (peak latency and 50% area latency).

Figure 5 shows the resulting SME values, averaged across participants, for each combination of scoring method and ERP component. Exact values are provided in supplemental Table S1, and RMS values are provided in supplemental Figure S1 and Table S1.

**Figure 5.**
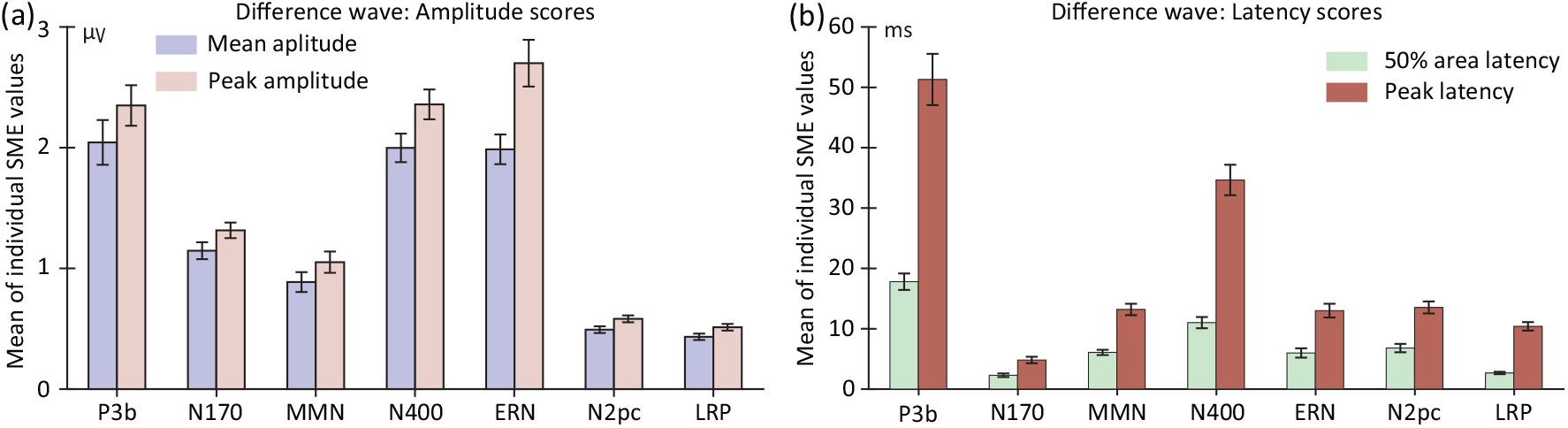
Standardized measurement error (SME) values, averaged across participants, for scores obtained from the difference waves used to isolate each of the seven ERP components. SME values are shown for the mean amplitude and peak amplitude scores (a) and for the peak latency score and 50% area latency score (b). Values were obtained using the time windows and electrode sites shown in Table 1. Error bars show the standard error of the mean across participants.

As was observed for the parent waves, the SME values for the difference waves were worse (larger) for peak amplitude than for mean amplitude. More precisely, the SME values were 1.20 times as large (i.e., 20% larger) for peak amplitude than for mean amplitude when averaged across the seven ERP components. Similarly, the SME values were worse for the peak latency measure than for the 50% area latency measure. Indeed, averaged across components, the SME values were 2.68 times larger for peak latency than for 50% area latency. This finding is consistent with the claim that 50% area latency is substantially more robust against noise than is peak latency (Clayson et al., 2013; Luck, 2014). In addition, the SME values varied widely across the seven ERP components, approximately paralleling the differences in SME values observed for the parent waveforms (Figure 4 a and b).

To provide statistical support for these observations, we conducted two repeated-measures analyses of variance (ANOVAs) on the SME values, one for the amplitude values and one for the latency values. Each ANOVA had two factors: scoring method and ERP component. For amplitude scores, the SME values were significantly worse (higher) for the peak amplitude method than for the mean amplitude method, F(1, 38) = 247.67, p < .001. For latency scores, the SME values were significantly worse (higher) for the peak latency method than for the 50% area latency method, F(1, 38) = 293.38, p < .001. For both amplitude and latency scores, SME values varied significantly across ERP components (amplitude: F(6, 33) = 69.49, p < .001; latency: F(6, 33) = 84.41, p < .001). The interaction between scoring method and ERP component was also significant for both amplitude scores (F(6, 33) = 28.98, p < .001) and latency scores (F(6, 33) = 43.69, p < .001).

#### 3.1.3. Variations in data quality across participants for each component and scoring procedure

Unlike psychometric reliability metrics, which typically provide a single value for a group of participants and are strongly influenced by the range of values across the group, a group-independent SME value is obtained for each individual participant (see also Clayson et al. (2021)). Figure 6 shows the single-participant SME values for each component (assessed from the difference waves) for four scoring methods, and the range of SME values across participants are summarized as histograms in Figure 7. Exact values are provided in the online repository for this paper (https://doi.org/10.18115/D5JW4R). Figures 6 and 7 make it clear that the SME values varied greatly across individual participants, with SME values being 3–5 times greater for some participants than for others. Section 3.3.1 will examine whether these individual differences in SME are consistent across the different ERP components.

**Figure 6.**
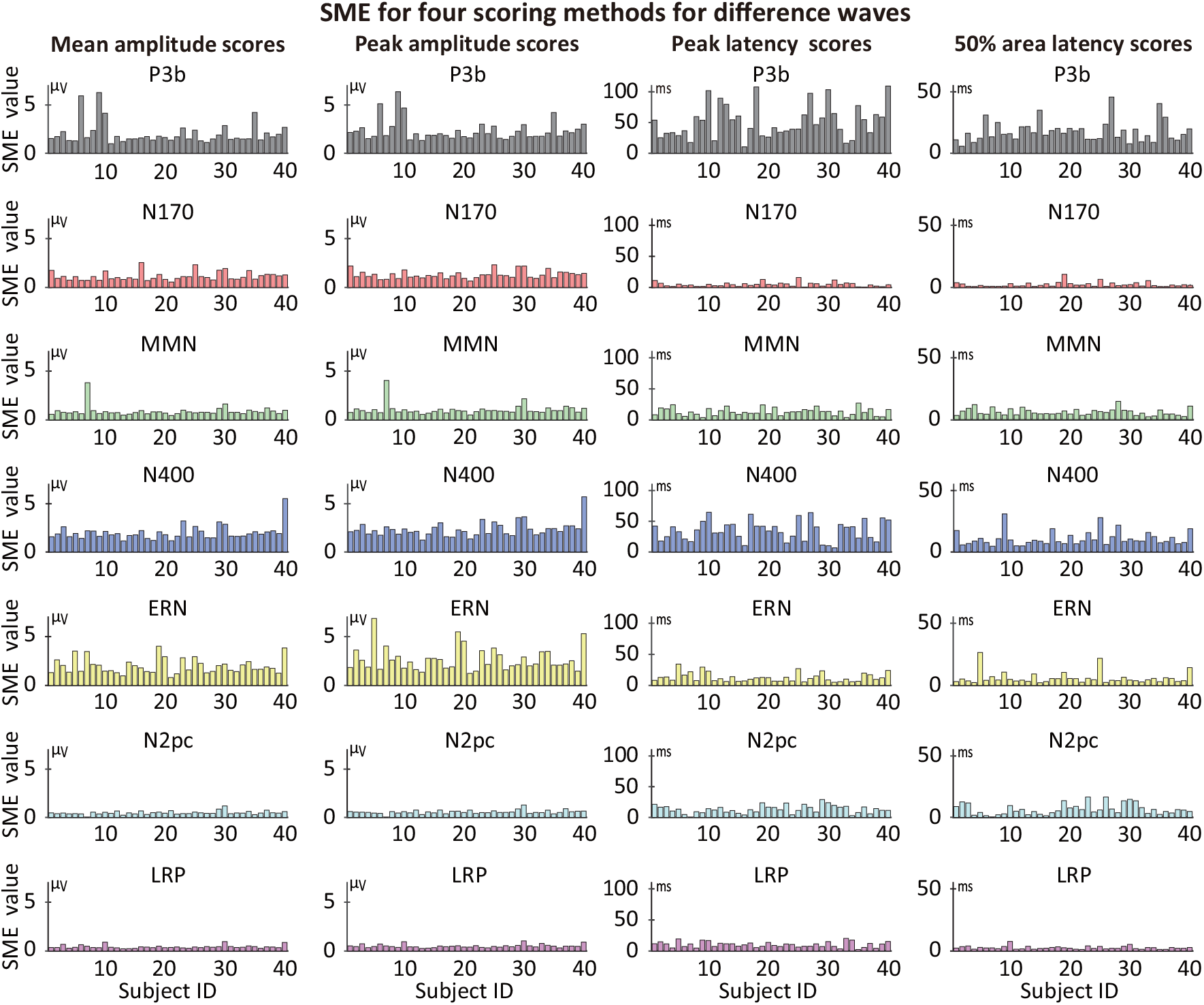
Single-participant SME values for four different scoring methods (mean amplitude, peak amplitude, peak latency, and 50% area latency), measured from difference waves for each of the seven ERP components. Each bar represents the SME value for one of the 40 participants.

**Figure 7.**
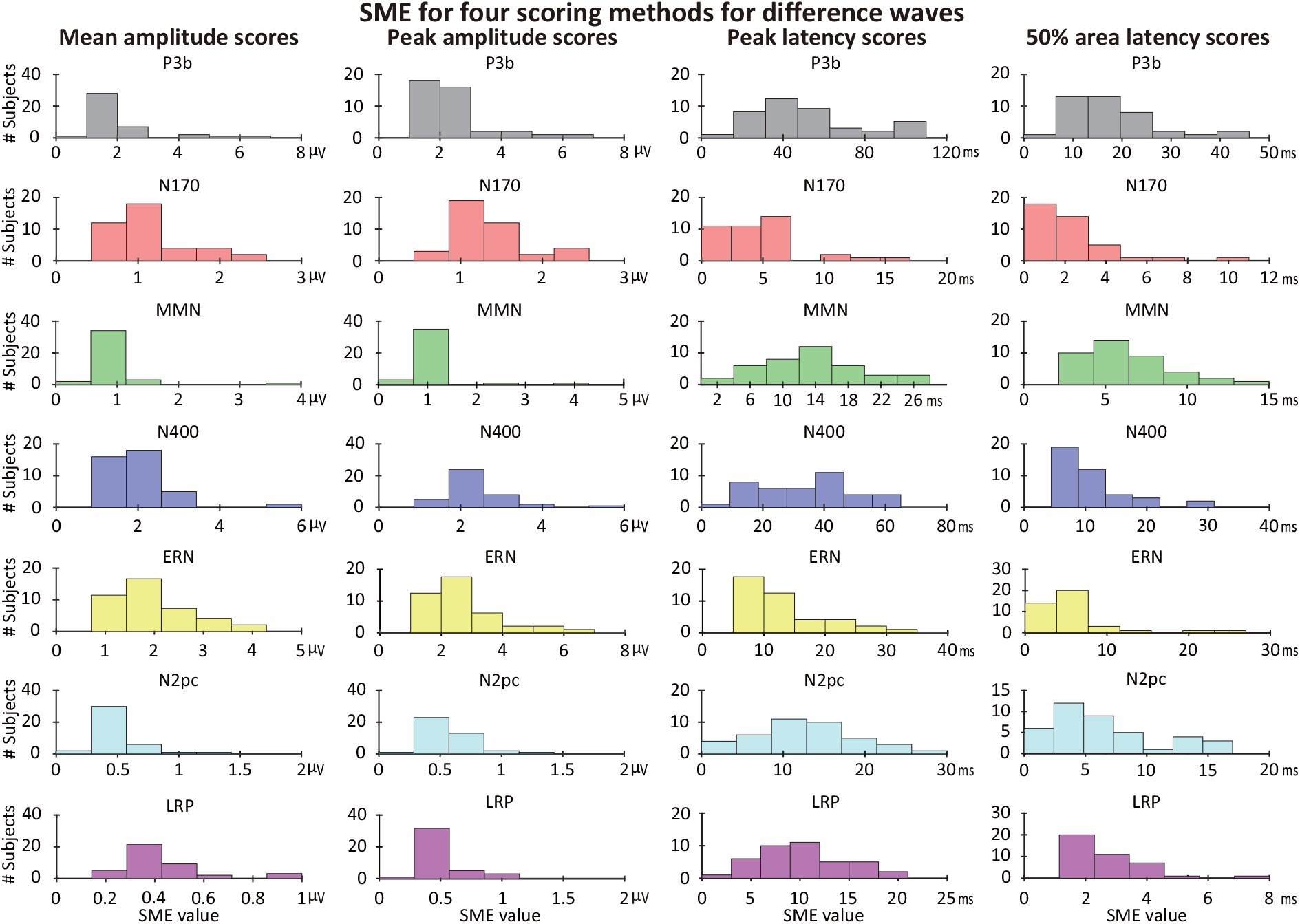
Histograms of single-participant standardized measurement error (SME) values for mean amplitude, peak amplitude, peak latency, and 50% area latency scores obtained from difference waves. For each component and each scoring method, the X axis was evenly divided into seven bins to reflect the different ranges of values for each plot.

Analogous single-participant plots and histograms are provided for the parent waveforms in supplementary Figures S2 and S3. The SME values for the parent waveforms also differed substantially across participants.

The SME values shown in Figures 4 - 7 and the associated supplementary materials provide an initial benchmark against which data quality from other data sets can be compared. That is, these values can be used as a comparison point to determine whether other recording environments or other variations on these paradigms lead to better or worse data quality. They also provide an initial benchmark for the variability in data quality across participants. To our knowledge, this is the first such set of data quality benchmark values across multiple ERP components and a reasonably large sample of participants. As noted in Section 1.3, however, these results may not generalize to other populations.

### 3.2. Variations in the number of trials and trial-to-trial EEG variability

Now that we have established that data quality varies substantially across paradigms, conditions, and participants, this section will examine two of the factors that are responsible for these differences, namely trial-to-trial EEG variability (quantified as the SD of the single-trial amplitudes) and the square root of the number of trials. For mean amplitude, Equation 1 states that the standard error of measurement for the mean amplitude score (which is the SME for mean amplitude) can be calculated for a given participant by simply dividing the SD by the square root of the number of trials. However, the extent to which the SD varies across paradigms, experimental conditions, and participants is an empirical question. In addition, the number of trials that remain after artifact rejection and the exclusion of trials with incorrect behavioral responses may vary across individuals. Thus, empirical data are needed to determine the extent to which SME values are actually influenced by the SD and the number of trials.

Figure 4 displays the mean across participants of the SME, the SD, and the square root of the number of trials for the parent waves corresponding to each of the seven ERP components. These values are shown for the mean amplitude scores on the left and for peak amplitude scores on the right. Equation 1 does not apply for peak amplitude, because the peak amplitude of an averaged ERP waveform is not equal to the average of the single-trial peak amplitudes, so empirical data are needed to determine how the SME for peak amplitude varies with the SD and the number of trials.

We focus on mean and peak amplitude scores in the following analyses because it is impossible to obtain meaningful single-trial latency scores and thereby estimate trial-to-trial variability for many of the components (e.g., the N2pc and LRP components, where the component is defined by a contralateral-minus-ipsilateral difference wave). In addition, we focus on the parent waves rather than the difference waves, because the number of trials varied across conditions for some of the components (e.g., the rare and frequent conditions in the P3b paradigm), and it is not clear how the number of trials for each condition should be combined when considering the SME for the difference between the conditions.

#### 3.2.1. The role of the number of trials in data quality for specific paradigms and conditions

We begin by considering the role of the number of trials per condition. As shown in Figure 4, when two conditions of a given paradigm differed in the number of trials, the SME for both mean amplitude and peak amplitude was worse (higher) in the condition with fewer trials. These differences were approximately linear with respect to the square root of the number of trials. For example, in the P3b and MMN paradigms, there were four times as many trials in the frequent category as in the rare category, and therefore the square root of the number of trials was twice as great in the frequent category as in the rare category. Correspondingly, the SME was approximately twice as great for the rare stimulus category as for the frequent stimulus category in these paradigms. The SME was also much greater for the error trials than for the correct trials in the ERN analysis, in which error trials were 10-25% as frequent as correct trials. The differences in SME between the conditions were statistically significant for the P3b, MMN, N400, and ERN analyses (for both mean and peak amplitude), but not for any of the other analyses (see Table 3).

Differences in the number of trials also partially explained differences in data quality between the different ERP components. For example, the data quality was considerably better (lower) for LRP (200 trials per condition) and N2pc (160 trials per condition) components than for the N170 component (80 trials per condition). However, differences in the number of trials did not explain all of the differences in data quality. For example, the SME for faces in the N170 paradigm was nearly identical to the SME for deviant stimuli in the MMN paradigm (see Figure 4 a), but there were approximately 2.5 times as many deviant trials in the MMN paradigm as face trials in the N170 paradigm. As shown in the next section, differences among ERP components in trial-to-trial EEG variability were also responsible for this and some of the other differences in data quality.

#### 3.2.2. The role of trial-to-trial EEG variability in data quality for specific paradigms and conditions

The trial-to-trial EEG variability (quantified as the SD of the single-trial scores) is shown in Figure 4c for mean amplitude scores and Figure 4d for peak amplitude scores. Just like the SME, the SD was significantly worse (larger) for peak amplitude than for mean amplitude for each of the 14 combinations of condition and ERP component (see Table S3). In addition, the SD varied widely across the seven different ERP components. The SD was lowest for the N170 and N2pc components, and substantially higher for the P3b, MMN, N400, ERN, and LRP.

Whereas the SME was worse (larger) for conditions with fewer trials than for conditions with more trials, the SD for a given component did not differ significantly across these conditions after correction for multiple comparisons (see Table 3). Even if significant differences had been seen, they could have been the result of the fact that the standard equation for estimating the SD is slightly biased by the number of observations. That is, the SD tends to be slightly underestimated when the number of observations is lower even when the appropriate degrees of freedom are used (Gurland & Tripathi, 1971). By contrast, the standard approach for estimating the variance across trials is unbiased, so the variance rather than the SD can be compared across conditions when the number of trials varies across conditions (see Figure S4).

In the N400 experiment, the trial-to-trial variability (SD) was greater for trials in which the target word was semantically related to the prime word than when the target and prime were semantically unrelated (which was statistically significant for mean amplitude; see Table 3). A potential explanation is provided in Section 4.2.

#### 3.2.3. The role of the number of trials in data quality for individual participants

The previous sections considered how the number of trials and the SD of the single-trial scores are related to differences in SME values across the seven components and across the pairs of experimental conditions used to define these components. We now turn to the role of these factors in explaining differences in data quality among individual participants.

The number of trials that were averaged together varied across participants as a result of artifact rejection and as a result of behavioral errors (in those analyses in which trials with errors were excluded: P3b, N170, N400, N2pc, LRP, and ERN). The number of trials varied greatly across participant for some components (e.g., P3b and ERN) but was relatively consistent across participants for other components (MMN, N170). These differences across components largely reflect the fact that some paradigms led to quite a bit of subject-to-subject variability in behavioral accuracy. Spreadsheets with the number of included and excluded trials for each participant for each component are provided in the online repository for this paper (https://doi.org/10.18115/D5JW4R).

Figure 8 shows scatterplots of the relationship between SME and the square root of the number of trials for each participant in each paradigm, separately for mean amplitude and peak amplitude. In many cases, the SME declined linearly as the square root of the number of trials increased. After correction for multiple comparisons, however, this effect was statistically significant only for the ERN error trials, in which there was an especially broad spread across individuals in the number of trials (see statistics embedded in Figure 8). Thus, individual differences in the number of trials remaining after artifact rejection and exclusion of errors had a substantial impact on data quality only when the number of trials varied considerably across participants.

**Figure 8.**
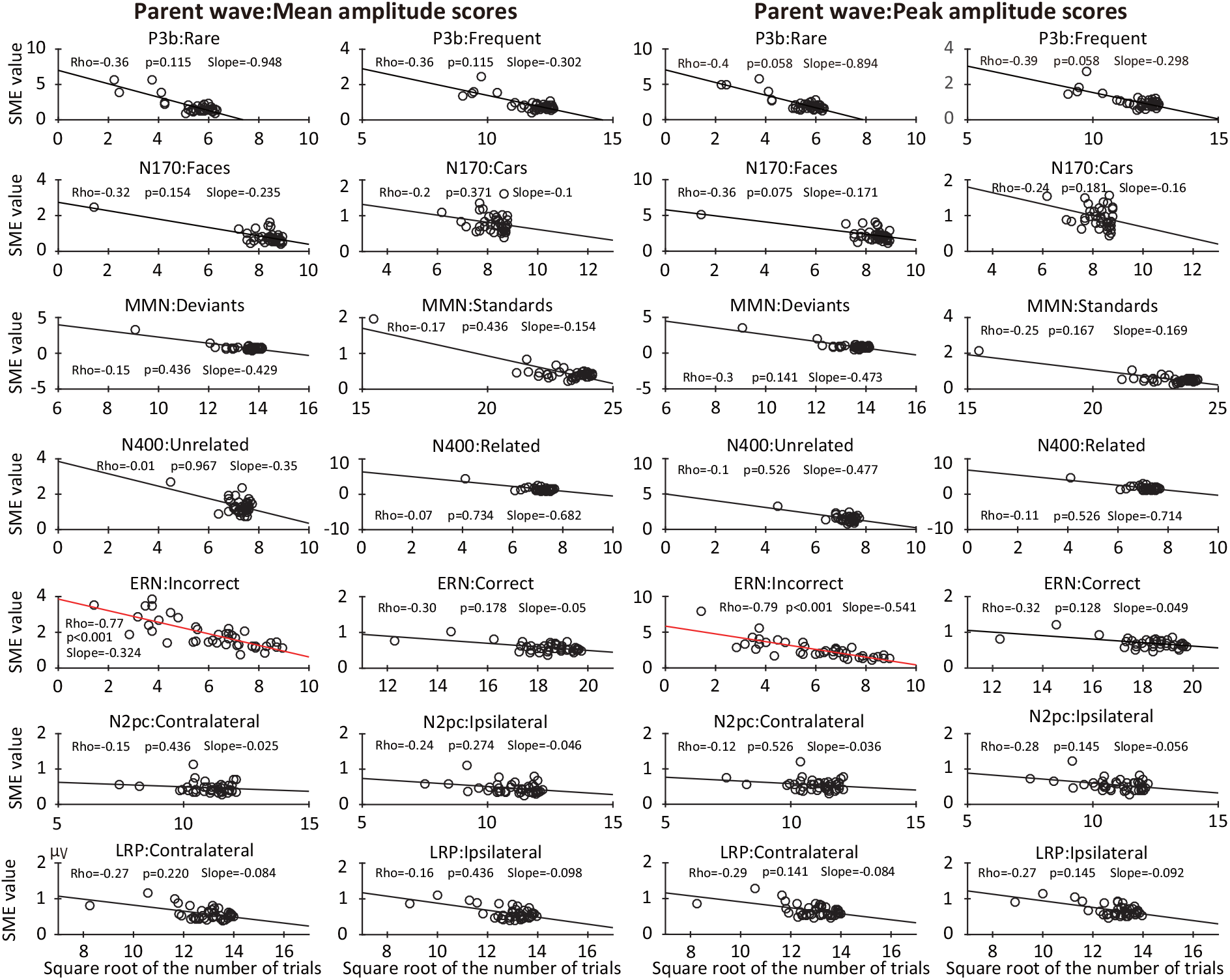
Scatterplots of the relationship between the standardized measurement error (SME, obtained from the parent waves) and the square root of the number of trials after rejection of trials with artifacts and behavioral errors. Scatterplots are shown for each of the seven components, separately for each of the two experimental conditions and for mean amplitude and peak amplitude scores. Each circle represents a single participant. The p values were corrected for multiple comparisons across the family of tests for each scoring method.

#### 3.2.4. The role of trial-to-trial EEG variability in data quality for individual participants

Differences between participants in EEG amplitude variability across trials played an important role in individual differences in SME. Figure 9 shows scatterplots of the relationship between SME and the single-trial SD obtained from parent waves for the mean amplitude and peak amplitude scores for each of the seven ERP components. All the cases showed a strong linear relationship, with correlations ranging from 0.79 to 0.99 for all cases except the error trials for the ERN component. In addition, with the exception of the ERN error trials, the correlations were substantially stronger for the SD (Figure 9) than for the square root of the number of trials (Figure 8). Thus, in the present data, individual differences in trial-to-trial EEG variability were the main driver of individual differences in data quality for the amplitude measures, although less so for the ERN (in which the number of trials varied considerably across participants). Note that the number of trials may be a more significant source of variation in SME in other paradigms or in other populations of research participants where the number of trials varies considerably across participants.

**Figure 9.**
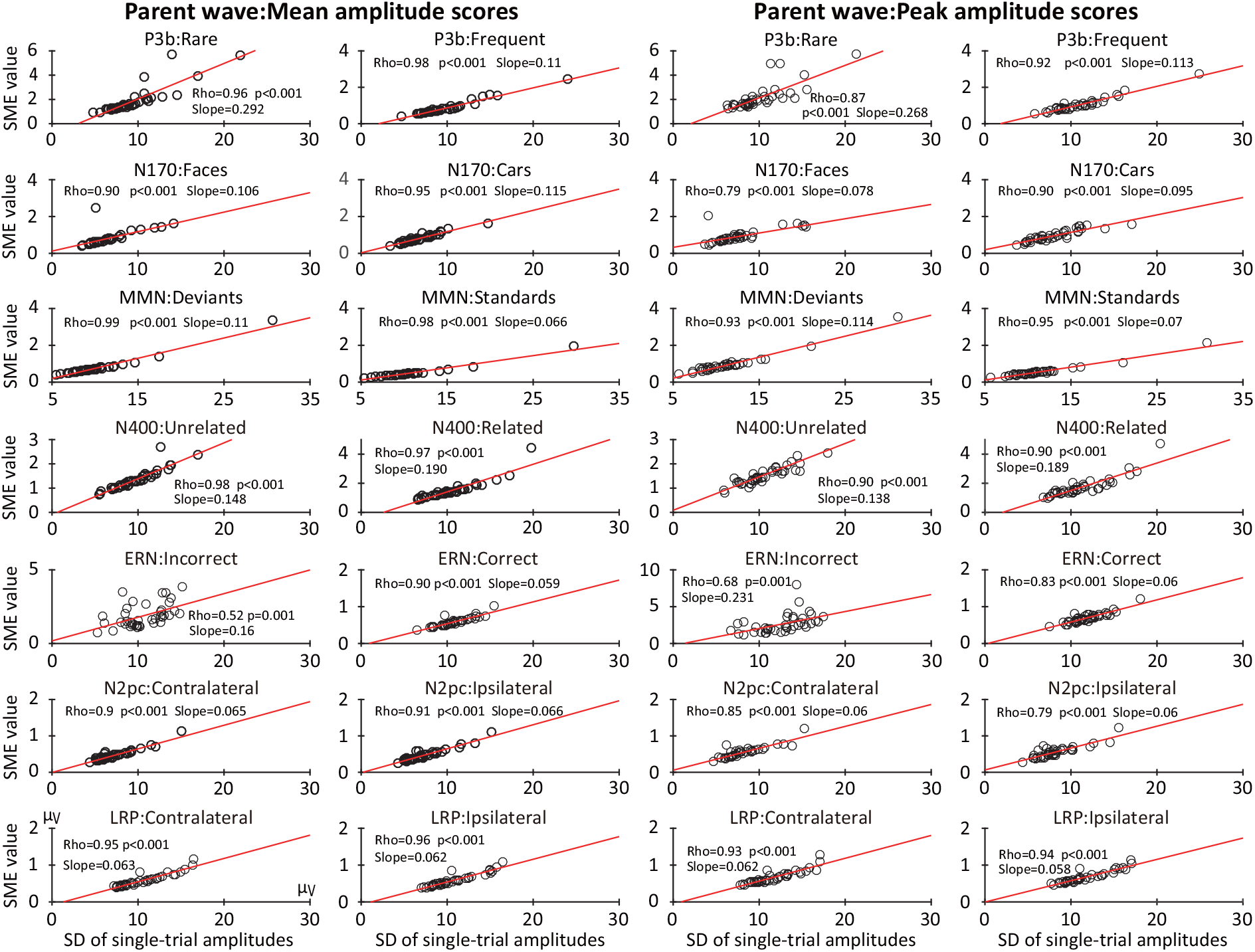
Scatterplots of the relationship between the standardized measurement error (SME, obtained from the parent waves) and the standard deviation (SD) of the single-trial scores for mean amplitude and peak amplitude scores. Scatterplots are shown for each of the seven components, separately for each of the two experimental conditions. Each circle represents a single participant. The p values were corrected for multiple comparisons across the family of tests for each scoring method.

Because of Equation 1, the SME for mean amplitude scores inevitably varies as a function of the SD. However, Equation 1 does not apply to peak amplitude scores, so the relationship between SD and SME for peak amplitude is an empirical question. Interestingly, we found that the SME for peak amplitude was strongly and approximately linearly related to the SD of the single-trial peak amplitudes in Figure 9. However, the correlations between SD and SME were slightly lower for peak amplitude than for mean amplitude. Also, given the lack of an analytic method for estimating the SME for peak amplitude scores, the strong and linear relationship between SD and SME for peak amplitude observed here may not hold for all experiments. Further, this relationship may not hold for other scoring methods. However, the fact that a strong linear relationship was observed for all seven ERP components examined here suggests that the variability in peak amplitude across trials is likely to be strongly associated with the SME for peak amplitude across a broad range of paradigms.

### 3.3. Are differences in data quality between participants consistent across paradigms and scoring procedures?

This section focuses on whether individual differences in data quality were consistent across the seven ERP components and the four scoring methods. That is, we asked whether the SME value for one participant in one paradigm (or one scoring method) predicts that individual’s SME in the other paradigms (or other scoring methods). Such a finding would indicate that some participants simply have poorer data quality than others. Alternatively, it is possible that the factors that determine data quality for one paradigm or scoring method are quite different from the relevant factors for other paradigms or scoring methods. For example, some scoring methods might be highly sensitive to high-frequency noise whereas other scoring methods might not. To distinguish between these possibilities, we examined the correlation between SME values across different paradigms and different scoring methods. We focused primarily on the bootstrapped SME values obtained from the difference waves, which could be validly assessed for all four scoring methods. We also examined how trial-to-trial variability (i.e., SD) correlated across the seven ERP components. Note that the SD was obtained only for amplitude measures, and only from the parent waves, because single-trial values are not defined for difference waves and are often impossible to obtain validly for latency scores.

#### 3.3.1. Consistency of SME across components (difference waves)

We first examined correlations in SME values across the seven ERP components. Figure 10 provides scatterplots and Spearman rank-order correlation values showing the relationship between the SME values for each pair of components, using the SME for the mean amplitude score (measured from difference waves). Analogous information is provided for the other scoring methods in Supplementary Figures S5, S6, and S7.

**Figure 10.**
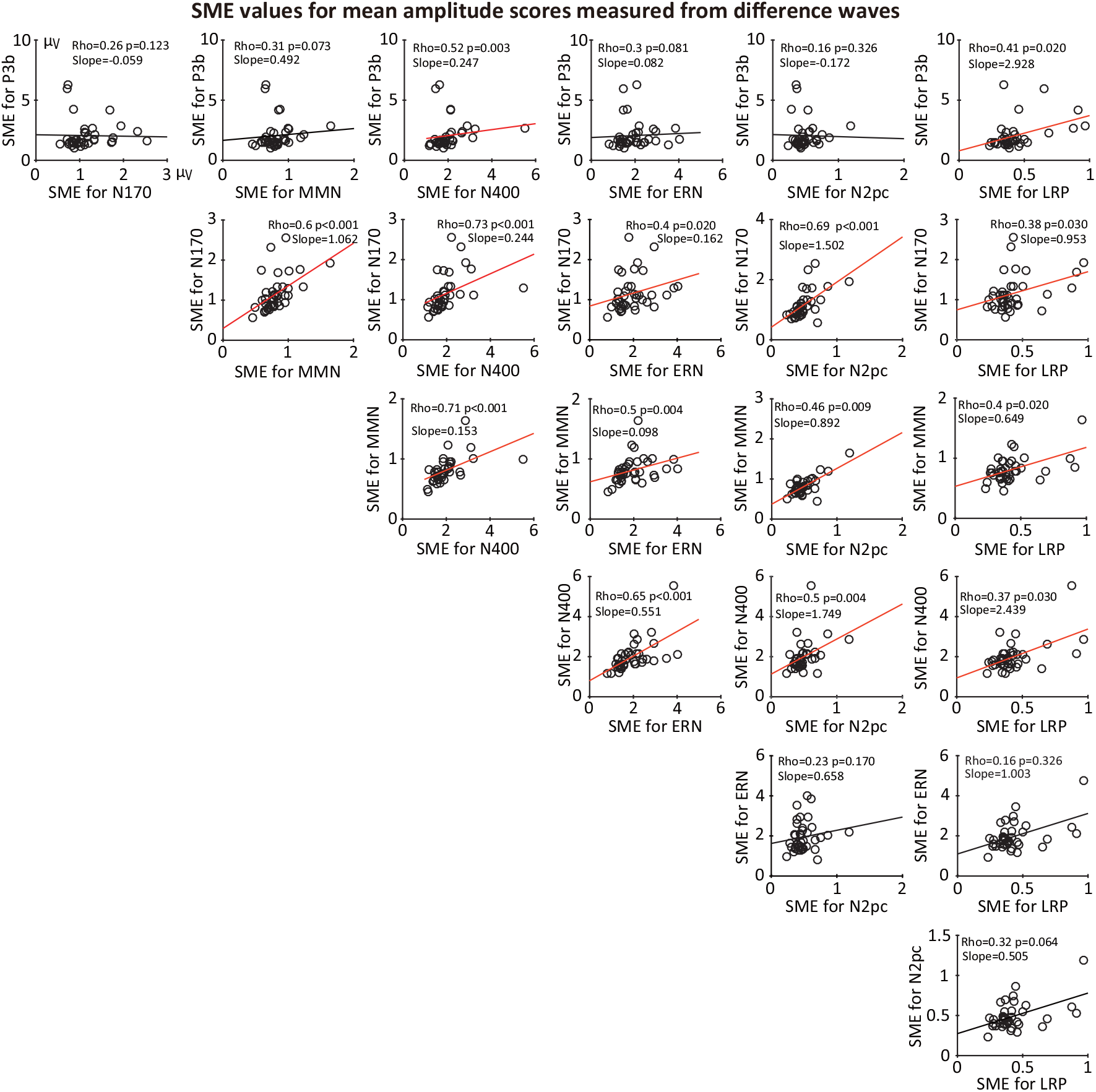
Scatterplots of the relationship between standardized measurement error (SME) values for each pair of ERP components (for mean amplitude scores obtained from difference waves). Each circle represents a single participant. The p values were corrected for multiple comparisons across this entire family of tests. To allow comparisons among the different components, Subject 7 was excluded from all scatterplots because the number of trials was zero after artifact rejection for one of the conditions of the N2pc experiment. Corresponding plots for peak amplitude, peak latency, and 50% area latency are provided in supplementary Figures S5, S6, and S7.

Significant positive correlations between SME values were observed for almost all pairs of components, indicating that a participant with poor data quality for one component tends to have poor data quality for other components as well. However, the correlations were far from perfect, and a few were not significant, suggesting that data quality is partially component-dependent. Similar results were obtained for the SME for peak amplitude scores (supplementary Figure S5). For the peak latency and 50% area latency scores, the SME values showed much weaker correlations across components (supplementary Figures S6 and S7). For these scoring methods, data quality appears to depend on different factors for different components. Supplementary Figures S8 and S9 show that similar results were obtained when we examined SD values rather than SME values.

To obtain an overall quantification of associations across components in SME values, as shown in Figure 11, we computed the intraclass correlation coefficient (ICC) for each scoring method. That is, we treated each component like a different rater of data quality for each participant and asked how consistently the different components “rated” the data quality. The SME values were first z-scored across participants for each component to put them into a consistent range of values. The ICC was 0.8 for the SME for mean amplitude score and 0.79 for the SME for peak amplitude score, indicating a reasonably high level of concordance of SME values across components for the amplitude measures. However, the ICC was only 0.35 for the SME for peak latency and 0.28 for the SME for 50% area latency, indicating a low level of concordance of SME values across components for the latency measures. These results confirm the observation that data quality was quite consistent across components for the amplitude scores but was much less consistent across components for the latency scores.

**Figure 11.**
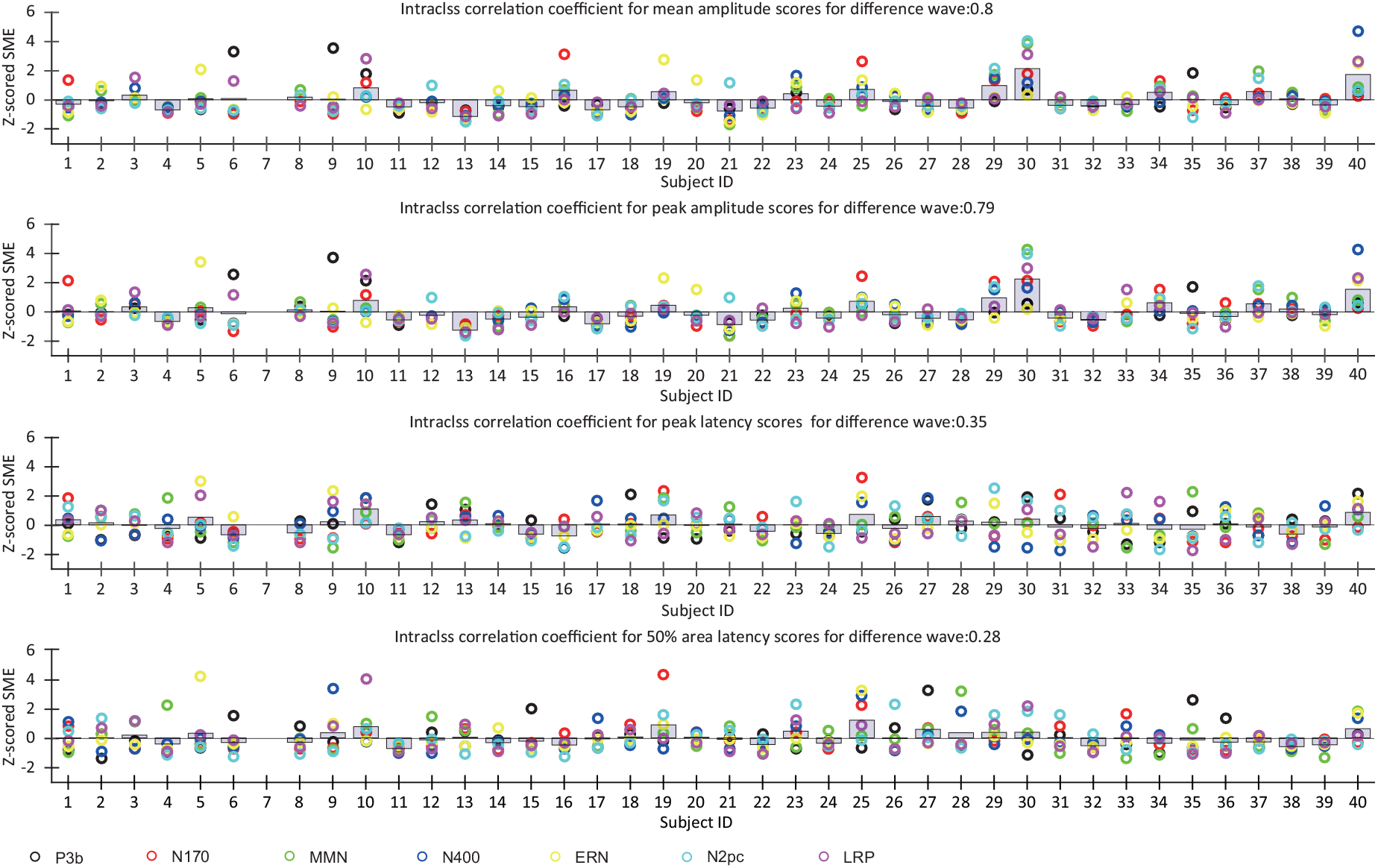
Variations in z-scored SME across participants for each component (obtained from difference waves).

#### 3.3.2. Consistency of SME across scoring methods (difference waves))

We next examined whether the SME was consistent across scoring methods for each component by asking whether the SME for a given scoring method was correlated with the SME for the other scoring methods, separately for each component. This was performed using scores obtained from the difference waves so that both amplitude and latency could be validly scored for every component. Figure 12 provides the resulting scatterplots and Spearman rank-order correlation values.

**Figure 12.**
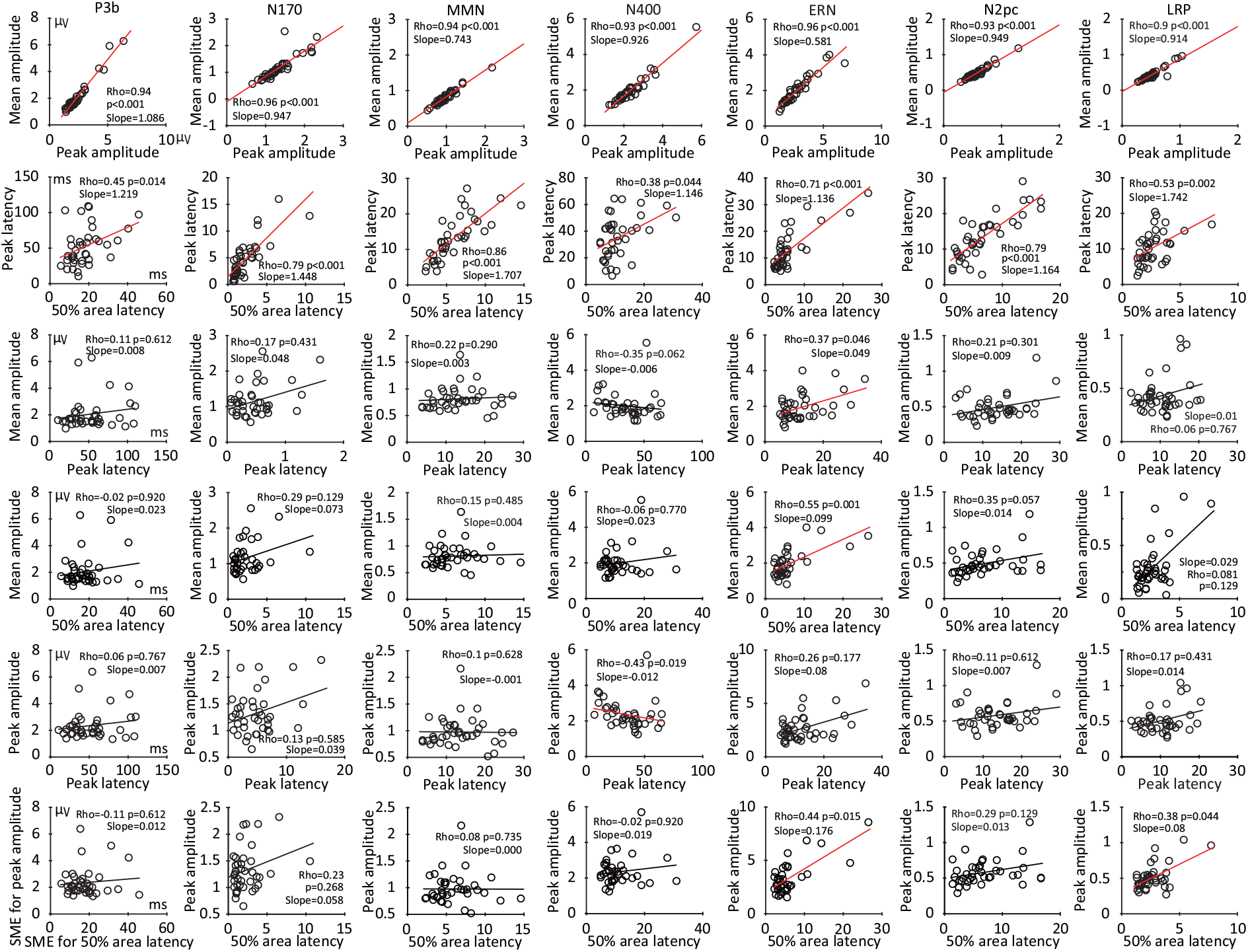
Scatterplots of the relationship between standardized measurement error (SME) values corresponding to each pair of the four scoring methods (mean amplitude, peak amplitude, peak latency, and 50% area latency), separately for each component. The SME values were obtained from the difference waves. The p values were corrected for multiple comparisons across this entire family of tests.

The SME values between mean and peak amplitude scoring methods were strongly correlated with each other for all seven components (Figure 12, top row). The SME values for 50% area latency and peak latency scores were also correlated with each other for all seven components (Figure 12, second row), but these correlations were not as strong as those between the SME values for the two amplitude measures. With a few exceptions, the SME for a given amplitude score and the SME for a given latency score were typically poorly correlated (Figure 12, rows 3-6). These results suggest that the factors that determine an individual participant’s data quality for an amplitude measure are often quite different from the factors that determine that individual’s data quality for a latency measure. In other words, “data quality” is not a single factor that is the same across amplitude and latency measures.

## 4. Discussion

In this study, we used the standardized measurement error (SME) to quantify the data quality across seven commonly studied ERP components, 40 individual participants, and four different scoring procedures (mean amplitude, peak amplitude, peak latency, and 50% area latency). We provided the SME values in multiple formats (Section 3.1) to allow other investigators to easily compare the SME values obtained here to their own SME values (which can be calculated using version 8 or later of ERPLAB Toolbox; Lopez-Calderon & Luck (2014)). All the scripts and results from the present study are available a folder named SME in the online repository for the ERP CORE (https://doi.org/10.18115/D5JW4R).

The present SME values can serve as a reference point for comparing data quality with different laboratories, different versions of the experimental paradigms, different participant populations, different recording systems, and different processing and analysis procedures. For example, a new laboratory could run one or more of these paradigms to determine whether they are obtaining comparable levels of data quality. Similarly, an established laboratory could determine whether their version of a given paradigm leads to better data quality (in which case the field could consider moving toward their methods) or worse data quality (in which case the laboratory could consider modifying their methods).

It is important to note that SME values could vary from those reported here solely as a result of differences in the number of trials and not because of differences in single-trial noise. The single-trial SD values provided here can be used to compare noise levels independent of trials. Alternatively, the trials a given dataset can be subsampled before computing the SME to provide a comparison with another dataset containing fewer trials.

### 4.1. Variations in data quality across paradigms, participants, and scoring procedures

The present study also found several interesting patterns of variation across paradigms, participants, and scoring procedures. First, data quality was somewhat better for mean amplitude scores than for peak amplitude scores, and much better for 50% area latency scores than for peak latency scores. These findings are consistent with previous studies using other approaches to assessing data quality. For example, Clayson et al. (2013) created a simulated noise-free ERP waveform and added simulated noise to examine the noise sensitivity of different scoring methods. Performance was quantified as the RMS error of the measurements across simulations relative to the noise-free data. Mean amplitude was less sensitive to noise than peak amplitude, and 50% area latency was less sensitive to noise than peak latency. Luck (2005) also found reduced variability for 50% area latency scores relative to peak latency scores using simulated data. The present results demonstrate that these patterns are also found in real data and across a broad range of experimental paradigms. Thus, researchers who currently use peak amplitude and/or peak latency should consider using mean amplitude and 50% area latency instead. Note, however, that 50% area latency is valid only when a component has been isolated via a difference wave or when the component is very large (e.g., P3b and N400; see Luck (2014)).

Second, data quality was much better for some components/paradigms than for others. For example, the SME for mean amplitude was approximately four times greater for the P3b component than for the N2pc component, and the SME for peak latency was more than ten times greater for the P3b component than for the N170 component (see Figures 4 and 5 and supplementary Table S1). However, the differences in amplitude between conditions tended to be larger in the paradigms with poorer data quality, and these factors may balance each other. Indeed, the effect size (Cohen’s d) for the difference between conditions was quite large for all seven components (see Table 3 in Kappenman et al. (2021)).

Third, the range of SME values across participants was quite large. Specifically, the SME was typically 5–10 times larger for the worst participant than for the best participant in a given paradigm (see Figures 6 and 7, and also the spreadsheets available at https://doi.org/10.18115/D5JW4R). This is a surprisingly wide range given that the participants were all neurotypical young adults attending a highly selective university and therefore relatively homogeneous in factors such as age, cognitive ability, self-control, and ability to understand and follow instructions. An even broader range of SME values would be expected for more diverse populations.

Effect sizes and statistical power are related to squared SME values (see Luck et al. (2021), especially Equations 3 and 5), which means that the participants with high SME values have an exponential impact on the likelihood of obtaining statistical significance. It would therefore be worthwhile for methodology researchers to focus on approaches to improving data quality for the most extreme cases.

### 4.2. Causes of variations in data quality

For mean amplitude scores, Equation 1 entails that the SME increases linearly as a function of the SD of the single-trial amplitudes and decreases linearly as a function of the square root of the number of trials, with no other contributing factors. In the present study, some of the differences in data quality across paradigms and across conditions within a paradigm were mainly a result of differences in the square root of the number of trials. For example, the SME values for peak amplitude and mean amplitude scores were approximately twice as great for the rare conditions as for the frequent conditions in the P3b and MMN paradigms, reflecting the fact that there were approximately four times as many trials in the frequent conditions as in the rare conditions. Similarly, SME values tended to be lower for paradigms with more trials (e.g., the LRP paradigm, with 200 trials per condition) than for paradigms with fewer trials (e.g., the N170 component, with 80 trials per condition).

However, the number of trials did not fully explain differences in SME across conditions and paradigms. For example, there were approximately 2.5 times as many deviant trials in the MMN paradigm as face trials in the N170 paradigm, but the SME values for mean amplitude were nearly identical (see Figure 4a). In these cases, the differences were necessarily due to differences in trial-to-trial amplitude variability, because that is the only other factor that impacts the SME for mean amplitude in Equation 1.

Interestingly, the SME for mean amplitude in the N400 paradigm was greater for semantically related trials than for semantically unrelated trials, even though the number of trials was the same, because the SD of the single-trial amplitudes was greater for the related trials than for the unrelated trials. This was unexpected, because the variability of a signal ordinarily increases with the magnitude of the signal (Brandmaier et al., 2018), and the N400 was much larger for the unrelated trials. This difference may reflect the fact that the association strength between the preceding prime word and the target word was much more variable for the related target words (association strength = 0.73 to 0.94) than for the unrelated target words (association strength = 0.00 to 0.01) (see Kappenman et al. (2021) for details). This may have led to greater variability N400 amplitude variability for the unrelated trials, creating a larger SD and SME.

One might also expect that trial-to-trial variability would be greater for the rare category than for the frequent category in an oddball paradigm, but this was not observed for either the P3b or the MMN (but see supplementary Figure S4 for an important caveat about comparing SD values for conditions that differ in the number of trials). These findings indicate the value of actually quantifying the trial-to-trial variability.

Note that trial-to-trial variability in cognitive processing can be theoretically important (Tamm et al., 2012; Ratcliff & McKoon, 2008). The SME would not be a good way to quantify neural variability, because it depends on the number of trials as well as the trial-to-trial variability. The SD is better because it is less dependent on the number of trials. However, the SD is influenced by nonneural sources of variability as well as neural sources. For example, differences in movement artifacts and skin potentials between groups or conditions could cause differences in SD between groups or conditions. The SD would be a useful way of comparing neural variability only if it was clear that nonneural sources of variability were unlikely to differ across groups or conditions. In addition, the true SD is underestimated by the sample SD when the number of trials becomes small (but there is a correction for this; see Gurland & Tripathi (1971)).

We also examined how the number of trials and the trial-to-trial EEG variability were related to the differences in SME across participants. The number of error trials varied considerably across participants for the ERN paradigm, and the SME for error trials was strongly and linearly related to the square root of the number of trials (Figure 8). Weaker and nonsignificant effects were seen for other components, in which the number of trials did not vary as much across participants. For all seven ERP components, however, differences among participants in SME were strongly predicted by individual differences in trial-to-trial EEG variability (Figure 9).

Unfortunately, these analyses were limited to amplitude measures obtained from the parent waves, because it was not straightforward to assess trial-to-trial variability for difference waves, and it was impossible to obtain valid latency measures from the parent waves for several of the components. In addition, Equation 1 is valid only for mean amplitude scores, so the factors that impact data quality for other scoring procedures cannot be determined analytically. The present analyses suggest that the effects of trial-to-trial variability and the number of trials are similar for peak amplitude and mean amplitude (compare the left and right halves of Figures 8 and 9). However, other factors may also play a role, especially for latency scores. For example, Luck (2014) speculated that peak latency will be difficult to measure precisely from a broad, low-amplitude waveform. Additional research will be needed to determine the factors that contribute to the SME for scoring methods such as peak latency and 50% area latency.

The question of why the SME varies across paradigms and participants can also be asked in terms of the types of signals and noise that are present in the EEG. For example, what is the relative impact of alpha-band EEG oscillations, low-frequency skin potentials, or line noise for a given amplitude or latency score? This will be an important topic for future research.

### 4.3. Consistency of data quality across paradigms and scoring procedures for individual participants

When EEG data are viewed in real time during a recording session, it sometimes seems obvious that the data from the current session are “clean” or “noisy”. This assumes that the data quality for a given participant will be “good” or “bad” on the basis of the raw EEG alone, independent of how the data are scored. As discussed in Section 1.2, however, the concept of data quality in ERP research must be defined with respect to the specific scores that will be obtained from the averaged ERP waveforms. Consequently, it is quite possible that a given participant could have “good” data quality for one component or scoring method but have “bad” data quality for another component or scoring method. However, it is also theoretically possible that individual differences in the raw EEG signal are the main driver of individual differences in data quality, with relatively little effect of the experimental paradigm or scoring method.

We addressed this issue by determining how the SME values were correlated across components. For the amplitude scores, we found significant correlations between the SME values for many pairs of components (see Figure 10 and supplementary Figure S5), and the intraclass correlation coefficients were fairly high (0.80 for mean amplitude and 0.79 for peak amplitude). However, the correlations in SME between pairs of components were quite weak for the two latency measures, with low intraclass correlation coefficients (0.35 for peak latency and 0.28 for 50% area latency).

We also asked whether the data quality for a given scoring procedure was correlated with the data quality for the other scoring procedures (see Figure 12). The SME values for the two amplitude scores were nearly perfectly correlated with each other for all seven components, but the SME values for the two latency scores were only modestly correlated with each other for most components. In addition, the correlations were quite low between the amplitude and latency SME values for most components.

Together, this pattern of correlations suggests that an individual’s ERP data quality is not purely a function of how “noisy” the EEG waveforms are. Instead, data quality is strongly impacted by whether the data are scored for amplitude or for latency and by which latency scoring procedure is used. Moreover, when latencies were scored, the data quality for one component was not a good predictor of data quality for most other components. However, when amplitudes were scored, data quality was highly consistent across components (although this may not be true when the different components are recorded in different sessions). These results reinforce the idea that ERP data quality depend on both the properties of a participant’s EEG signal and the scoring method.

### 4.4. Concluding comments

Although data quality is obviously important in ERP research, we know of no prior efforts to systematically quantify ERP data quality across a large number of paradigms, participants, and scoring procedures. The present results indicate that data quality varies quite widely across these variables. We hope that this study inspires other researchers to quantify their data quality, which is an important first step toward increasing the quality of the data and therefore the statistical power of their experiments. Toward that end, we have made it trivial to compute the SME for mean amplitude in ERPLAB Toolbox (version 8 and later), and we have provided example scripts for using bootstrapping to compute the SME for peak amplitude, peak latency, and 50% area latency (https://doi.org/10.18115/D5JW4R).

SME values can be very helpful in performing power analyses. In particular, because the SME varies linearly with the square root of the number of trials, it is possible to predict how the SME will change if the number of trials is increased or decreased for a given experiment. Our original SME paper (Luck et al., 2021; Baker et al., 2021) provides a detailed description of how SME scores can be used to estimate effect sizes, which can then be used in power analyses. In addition, it describes how to convert SME values into measurement error variance, which can in turn be plugged into a power calculator that predicts how power will vary according to any combination of number of participants and number of trials (Baker et al., 2021).

It would be very helpful for researchers to provide SME benchmarks for other paradigms, participant populations, and scoring procedures. For example, a recent study by Isbell & Grammer (2022) examined the SME for the ERN in children between the ages of 5 and 7 using a child-friendly version of the Go/NoGo task. They found that the SME values were substantially larger than those found for the ERN in the present study, which is not surprising given the challenges involved in recording the EEG from children.

Another useful direction for future research would be to assess the impact of different data processing pipelines. For example, when a strict artifact rejection threshold is imposed, this may reduce trial-to-trial variability and thereby improve the data quality, but it will also reduce the number of trials which may degrade the data quality. The SME can provide an objective criterion for determining which parameters or algorithms are optimal (see Luck (2022)). However, it will be important for such research to determine whether a given set of parameters or algorithms leads to a bias in the amplitude or latency scores, which would not be evident in the SME values but could lead to incorrect conclusions. Ideally, researchers should strive to obtain scores that are both accurate (have minimal bias) and precise (have low SME values).

## Declaration of interest

The authors declare no competing financial interests.

## Acknowledgments

This study was made possible by grant R01MH087450 from the National Institute of Mental Health to SJL.

## Appendix A Supplementary materials

**Figure S1.**
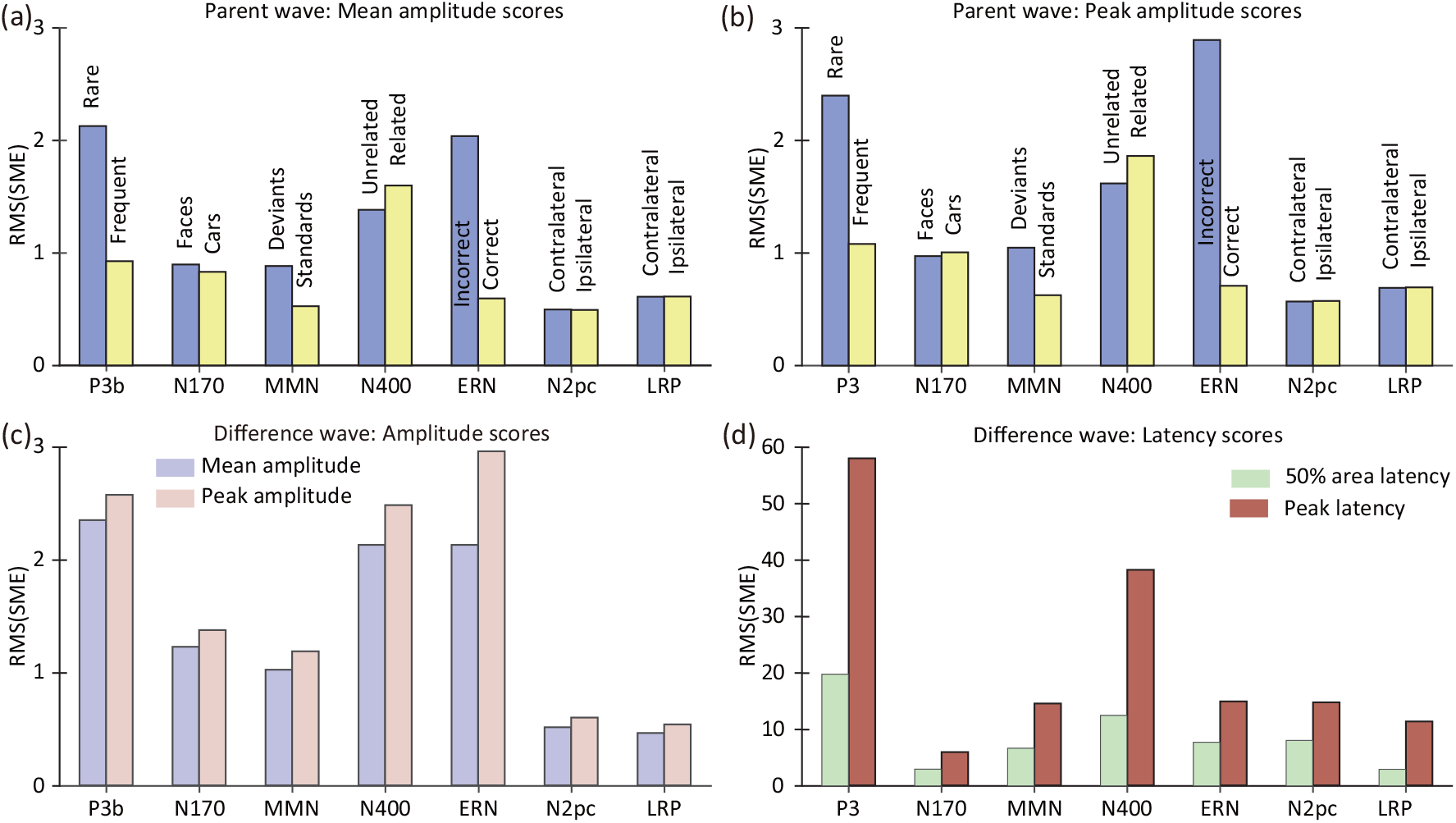
Root mean square of the single-participant SME values for the parent waves (a, b) and the difference waves (c, d). For the parent waves, the SME was computed only for mean amplitude (a) and peak amplitude (b), because latency values could not be validly measured from the parent waves in several cases. For the difference waves, the SME could be calculated for both the amplitude scores (c) and the latency scores (d).

**Table S1.** Mean (± standard error) of the average of SME values across participants, for each of the parent waves and for the difference waves for each of the seven ERP components.

| Parent wave | P3b |  | N170 |  | MMN |  | N400 |  |
| --- | --- | --- | --- | --- | --- | --- | --- | --- |
|  | Rare | Frequent | Faces | Cars | Deviants | Standards | Unrelated | Related |
| Mean amplitude | 1.83 ± 0.17 | 0.85± 0.06 | 0.81 ±0.06 | 0.79 ± 0.04 | 0.76± 0.07 | 0.45± 0.04 | 1.32 ±0.07 | 1.49±0.09 |
| Peak amplitude | 2.18± 0.16 | 1.01± 0.06 | 0.91 ±0.05 | 0.96 ±0.05 | 0.93 ± 0.08 | 0.55± 0.05 | 1.56±0.07 | 1.74 ± 0.11 |
| Parent wave | ERN |  | N2pc |  | LRP |  | – |  |
|  | Incorrect | Correct | Contralateral | Ipsilateral | Contralateral | Ipsilateral | – | – |
| Mean amplitude | 1.88 ±0.12 | 0.58± 0.02 | 0.47±0.03 | 0.47± 0.03 | 0.59 ±0.03 | 0.59 ±0.03 | – | – |
| Peak amplitude | 2.59 ±0.21 | 0.70± 0.02 | 0.55±0.03 | 0.55± 0.03 | 0.67 ±0.03 | 0.68±0.03 | – | – |
| Difference wave | P3b | N170 | MMN | N400 | ERN | N2pc | LRP | – |
| Mean amplitude | 2.04 ± 0.19 | 1.15± 0.07 | 0.89±0.08 | 2.00± 0.12 | 1.99 ±0.12 | 0.49±0.03 | 0.43±0.03 | – |
| Peak amplitude | 2.35 ±0.17 | 1.32± 0.06 | 1.05± 0.09 | 2.36 ±0.12 | 2.70±0.19 | 0.58±0.03 | 0.51±0.03 | – |
| 50% area latency | 17.85±1.37 | 2.30±0.30 | 6.09±0.44 | 11.04±0.94 | 6.00±0.78 | 6.82±0.70 | 2.68±0.20 | – |
| Peak latency | 51.46±4.26 | 4.83±0.54 | 13.24±0.95 | 34.75±2.53 | 13.04±1.15 | 13.56±1.00 | 10.42±0.71 | – |

**Table S2.**
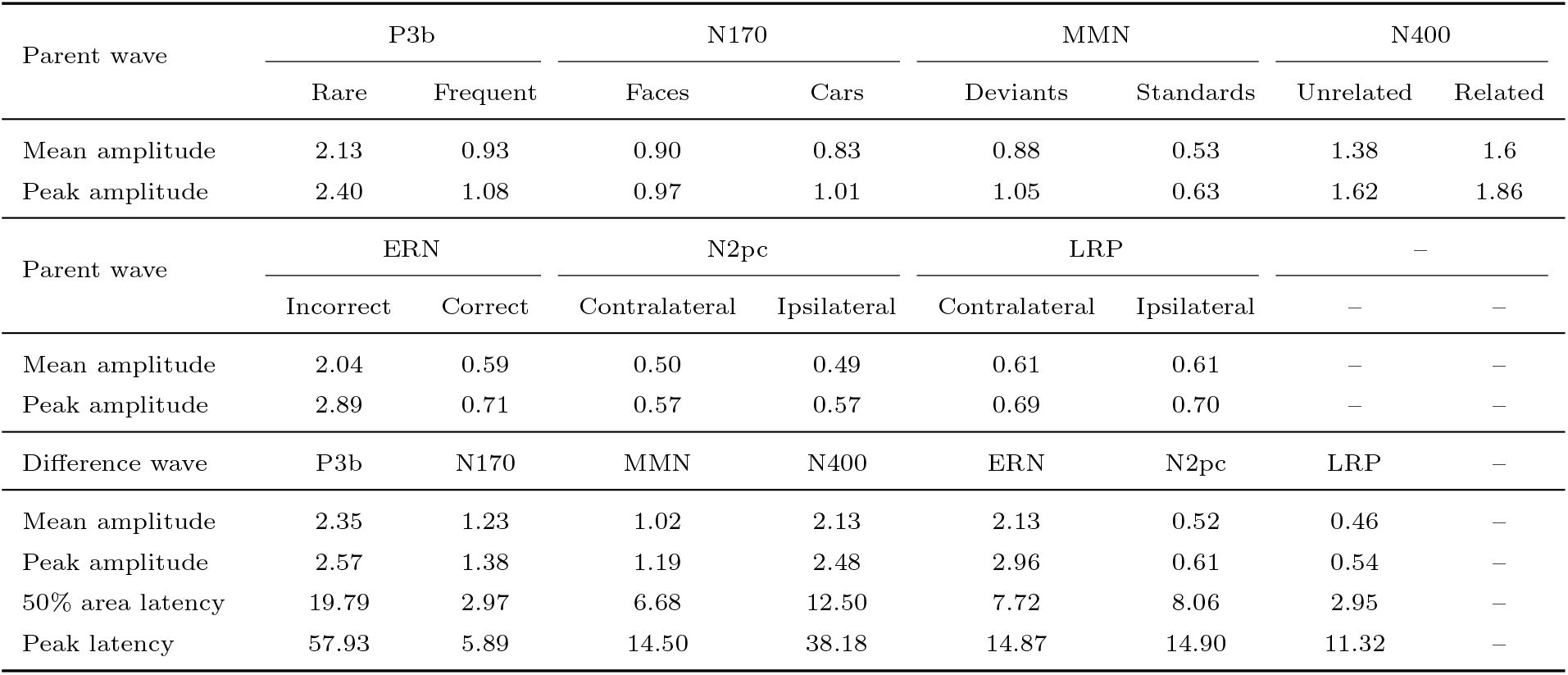
Root mean square (RMS) of SME values across participants, for each of the parent waves and for the difference waves for each of the seven ERP components.

**Table S3.** Paired t tests comparing SME or SD for the peak versus mean amplitude scoring methods for each ERP component, separately for each experimental condition (corrected for multiple comparisons across the family of 14 tests for each dependent variable). Note that the degrees of freedom were lower for the N2pc because Subject 7 was excluded from the N2pc analyses.

| Parent wave | P3b |  | N170 |  | MMN |  | N400 |  | ERN |  | N2pc |  | LRP |  |
| --- | --- | --- | --- | --- | --- | --- | --- | --- | --- | --- | --- | --- | --- | --- |
|  | t(39) | p | t(39) | p | t(39) | p | t(39) | p | t(39) | p | t(38) | p | t(39) | p |
| SME for condition 1 | -7.13 | <.001 | -4.89 | <.001 | -12.00 | <.001 | -8.94 | <.001 | -6.41 | <.001 | -10.32 | <.001 | -12.19 | <.001 |
| SME for condition 2 | -12.07 | <.001 | -9.35 | <.001 | -13.86 | <.001 | -7.97 | <.001 | -15.46 | <.001 | -11.08 | <.001 | -9.98 | <.001 |
| SD for condition 1 | -3.74 | 0.001 | -5.25 | <.001 | -10.60 | <.001 | -8.63 | <.001 | -7.49 | <.001 | -9.00 | <.001 | -7.66 | <.001 |
| SD for condition 2 | -8.33 | <.001 | -8.33 | <.001 | -13.51 | <.001 | -5.70 | <.001 | 13.60 | <.001 | -8.37 | <.001 | -7.48 | <.001 |

**Table S4.** Paired t tests comparing the variance in the single-trial amplitudes between the two experimental conditions for each ERP component, separately for mean amplitude and peak amplitude (corrected for multiple comparisons across the family of tests for each scoring method). Note that the degrees of freedom were lower for the N2pc because Subject 7 was excluded from the N2pc analyses.

| Parent wave | P3b |  | N170 |  | MMN |  | N400 |  | ERN |  | N2pc |  | LRP |  |
| --- | --- | --- | --- | --- | --- | --- | --- | --- | --- | --- | --- | --- | --- | --- |
|  | t(39) | p | t(39) | p | t(39) | p | t(39) | p | t(39) | p | t(38) | p | t(39) | p |
| Variance for mean amplitude | -1.64 | 0.382 | -0.14 | 0.904 | -0.57 | 0.799 | -2.75 | 0.063 | 0.12 | 0.904 | 0.63 | 0.799 | -0.91 | 0.799 |
| Variance for peak amplitude | -2.11 | 0.144 | -1.27 | 0.371 | -0.26 | 0.892 | -1.90 | 0.152 | 2.76 | 0.063 | 0.14 | 0.892 | -0.50 | 0.865 |

**Figure S2.**
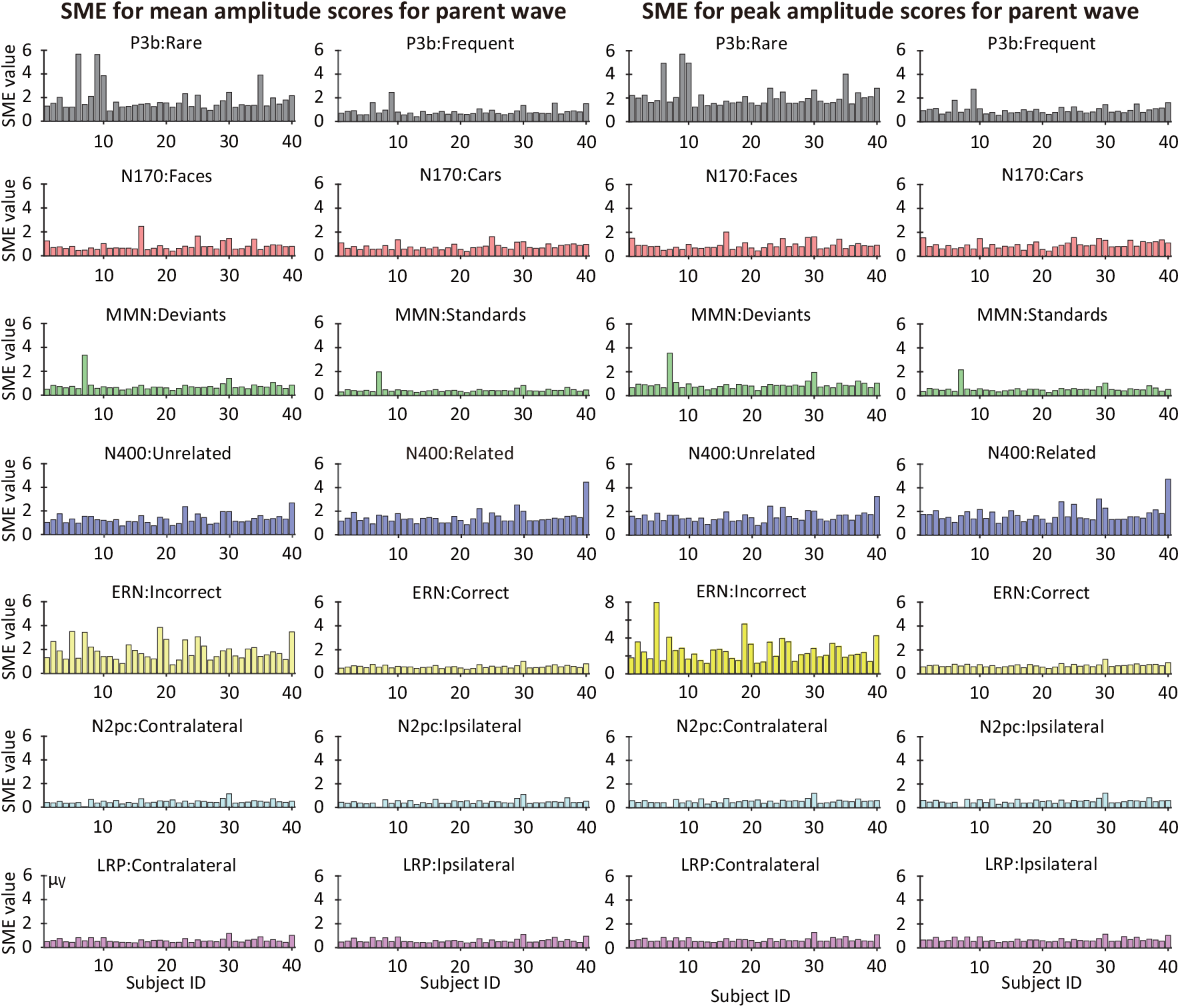
Single-participant SME values for mean amplitude and peak amplitude scores, quantified from the parent waves for each of the seven ERP components. Each bar represents one of the 40 participants. Values are provided only for mean amplitude and peak amplitude because latency values could not be validly measured from the parent waves in several cases.

**Figure S3.**
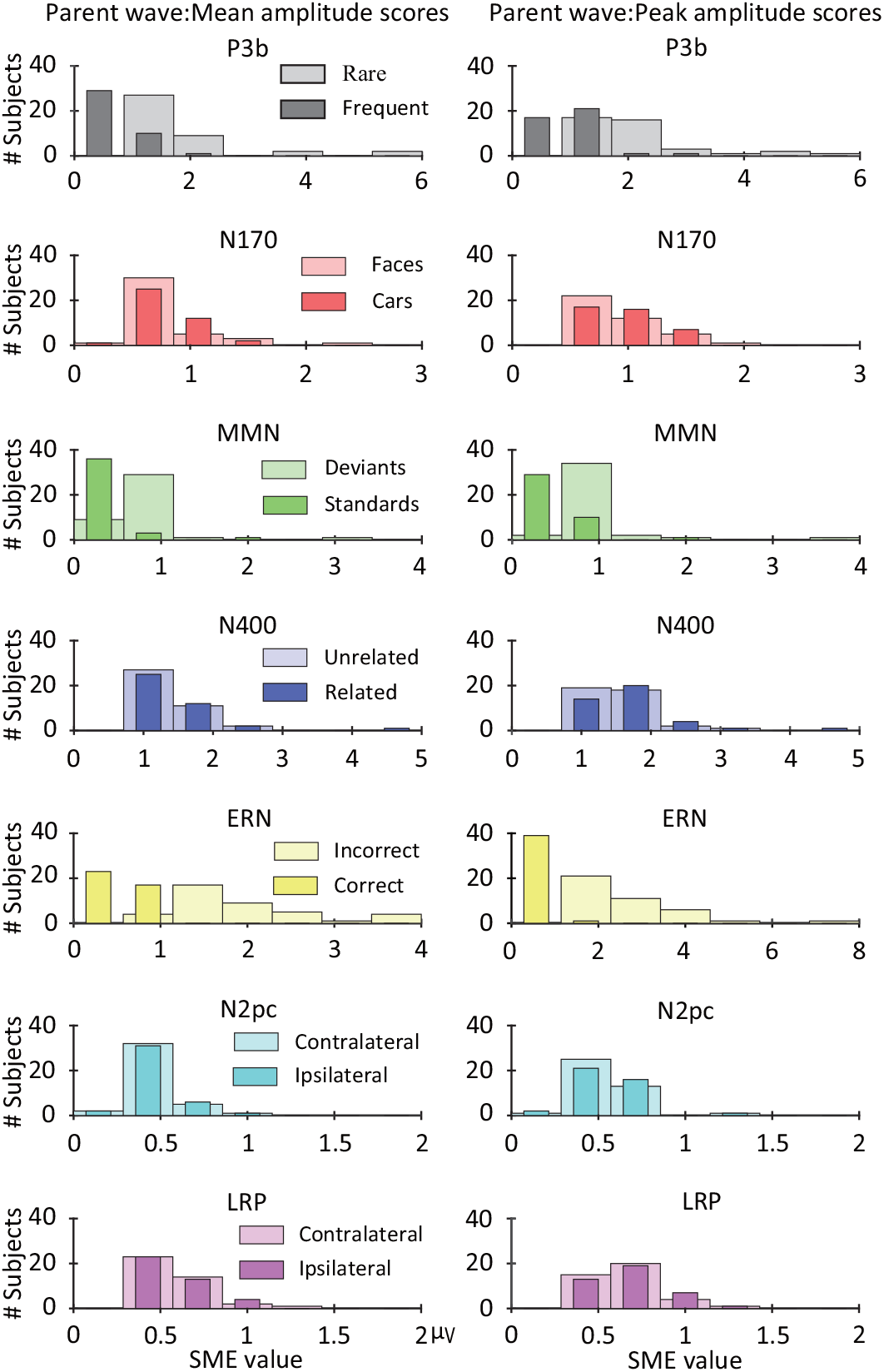
Histograms of single-participant SME values for mean amplitude and peak amplitude scores obtained from parent waves. For each component and each scoring method, the X axis was evenly divided into seven bins to reflect the different ranges of values for each plot. Values are provided only for mean amplitude and peak amplitude because latency values could not be validly measured from the parent waves in several cases.

**Figure S4.**
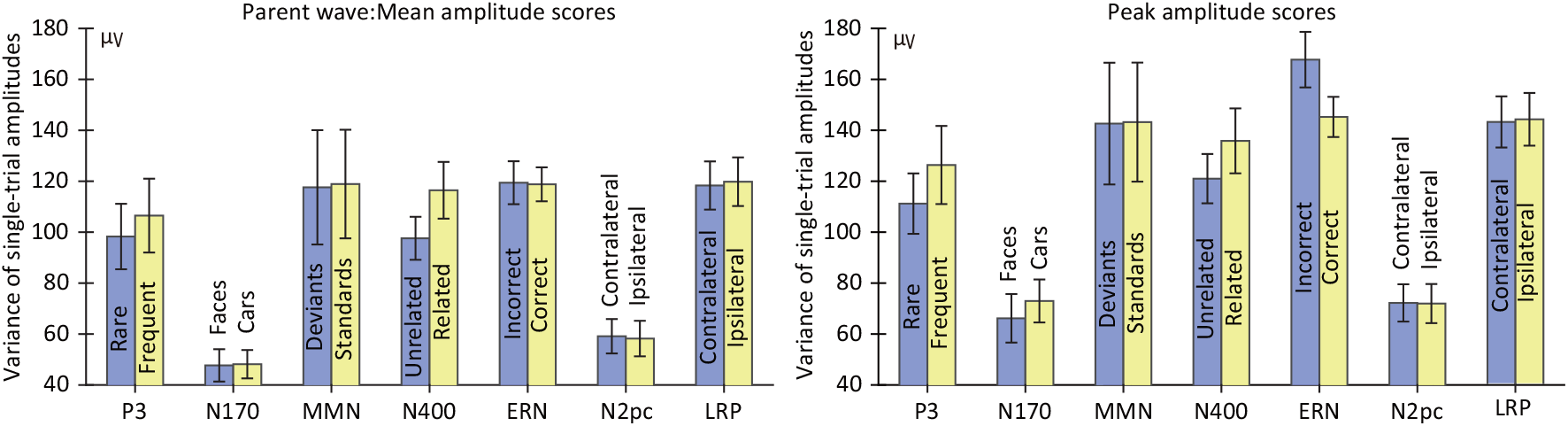
Analysis of the effects of variations in the number of trials on estimates of the variance in single-trial amplitude values. Panels c and d in Figure 4 of the main manuscript showed that the mean standard deviation (SD) across trials was numerically smaller in conditions with fewer trials than in conditions with more trials in several cases (for P3b mean and peak amplitude; for MMN mean and peak amplitude; for ERN mean amplitude). Although not statistically significant, these small differences in SD may reflect the fact that the equation typically used to estimate the SD is biased by the number of trials, underestimating (on average) the true SD by a progressively larger amount as the number of observations decreases (Gurland & Tripathi, 1971). As a result, the estimated SD for a condition with fewer trials will tend to be lower than the estimated SD for a condition with more trials, even if the true trial-to-trial variability is equivalent (all else being equal). The bias in the estimate of SD is a result of the square root operation used to convert the variance to the SD, and the equation for estimating the variance across trials is not biased by the number of trials. In the present figure, we show the variance rather than the SD across trials , averaged across participants for each condition. No significant differences between the rare and frequent conditions of the P3b, MMN, and ERP paradigms were present for the variance (see supplementary Table S4). Error bars show the standard error of the mean across participants.

**Figure S5.**
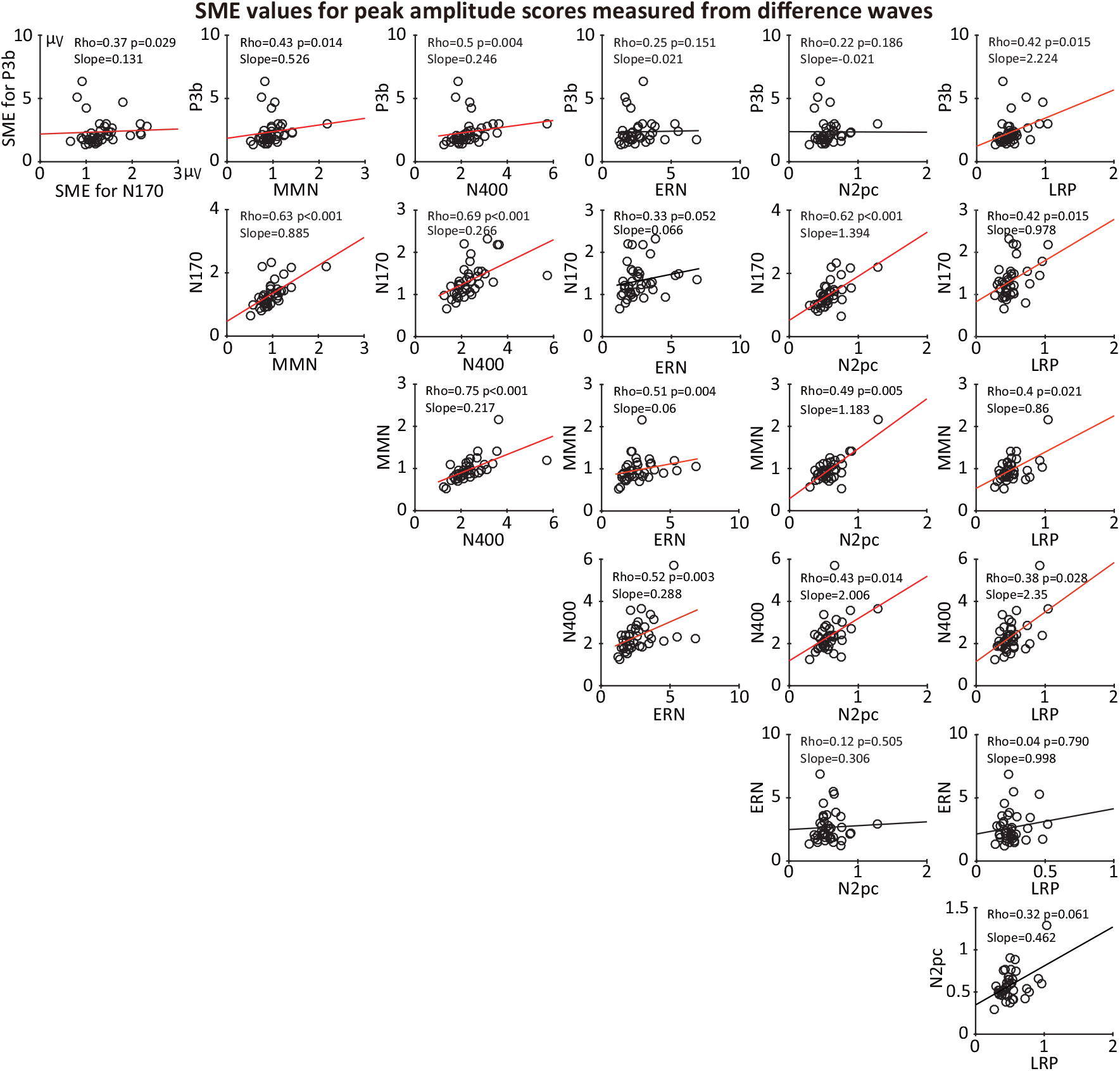
Scatterplots of the relationship between each pair of components for the SME values obtained for peak amplitude from the difference waves. Each circle represents a single participant. The p values were corrected for multiple comparisons across this entire family of tests. Note that Subject 7 was excluded from all correlations because the number of trials for one of conditions was zero for this participant in one condition of the N2pc paradigm.

**Figure S6.**
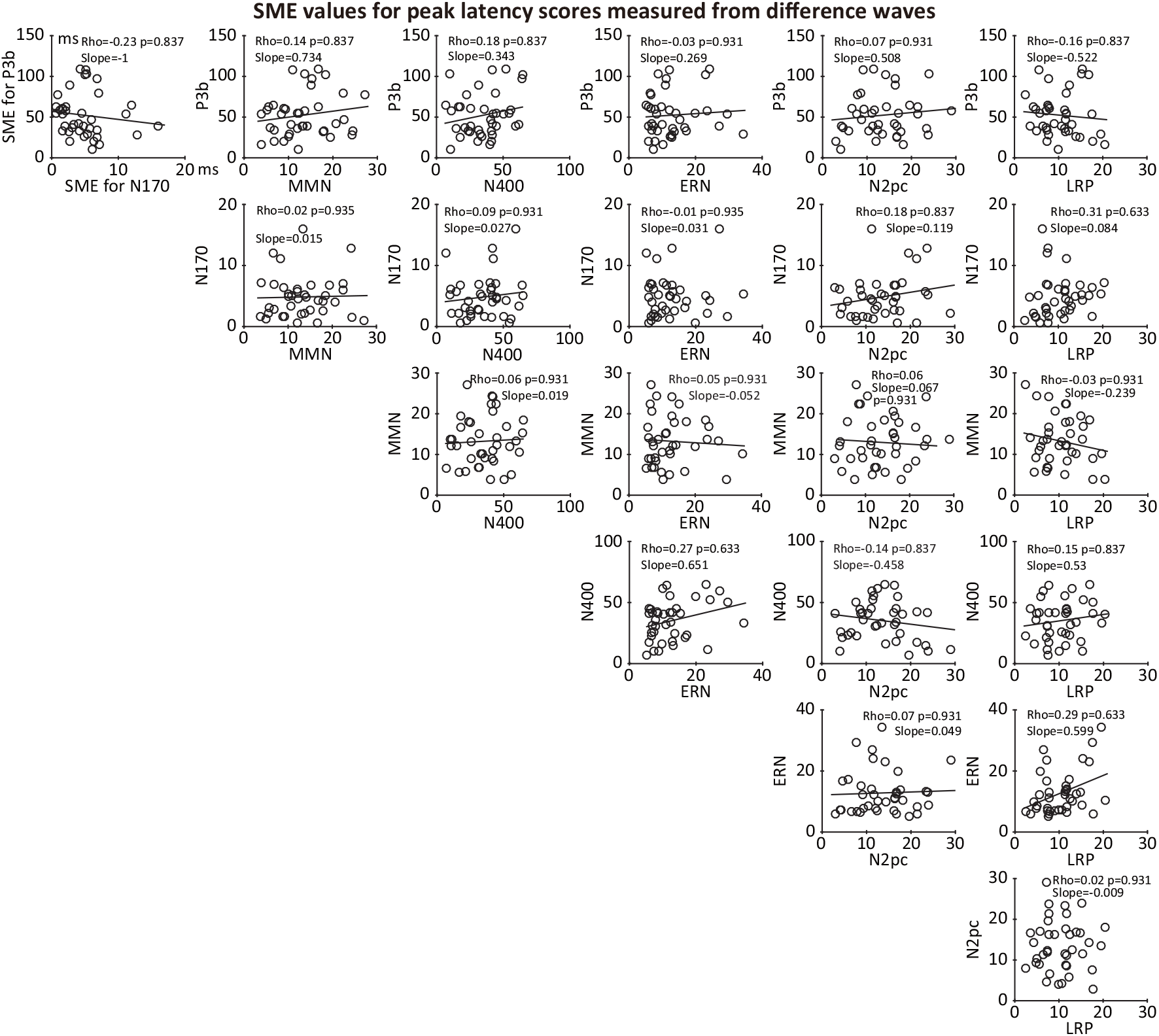
Scatterplots of the relationship between each pair of components for the SME values obtained for peak latency from the difference waves. Each circle represents a single participant. The p values were corrected for multiple comparisons across this entire family of tests. Note that Subject 7 was excluded from all correlations because the number of trials for one of conditions was zero for this participant in one condition of the N2pc paradigm.

**Figure S7.**
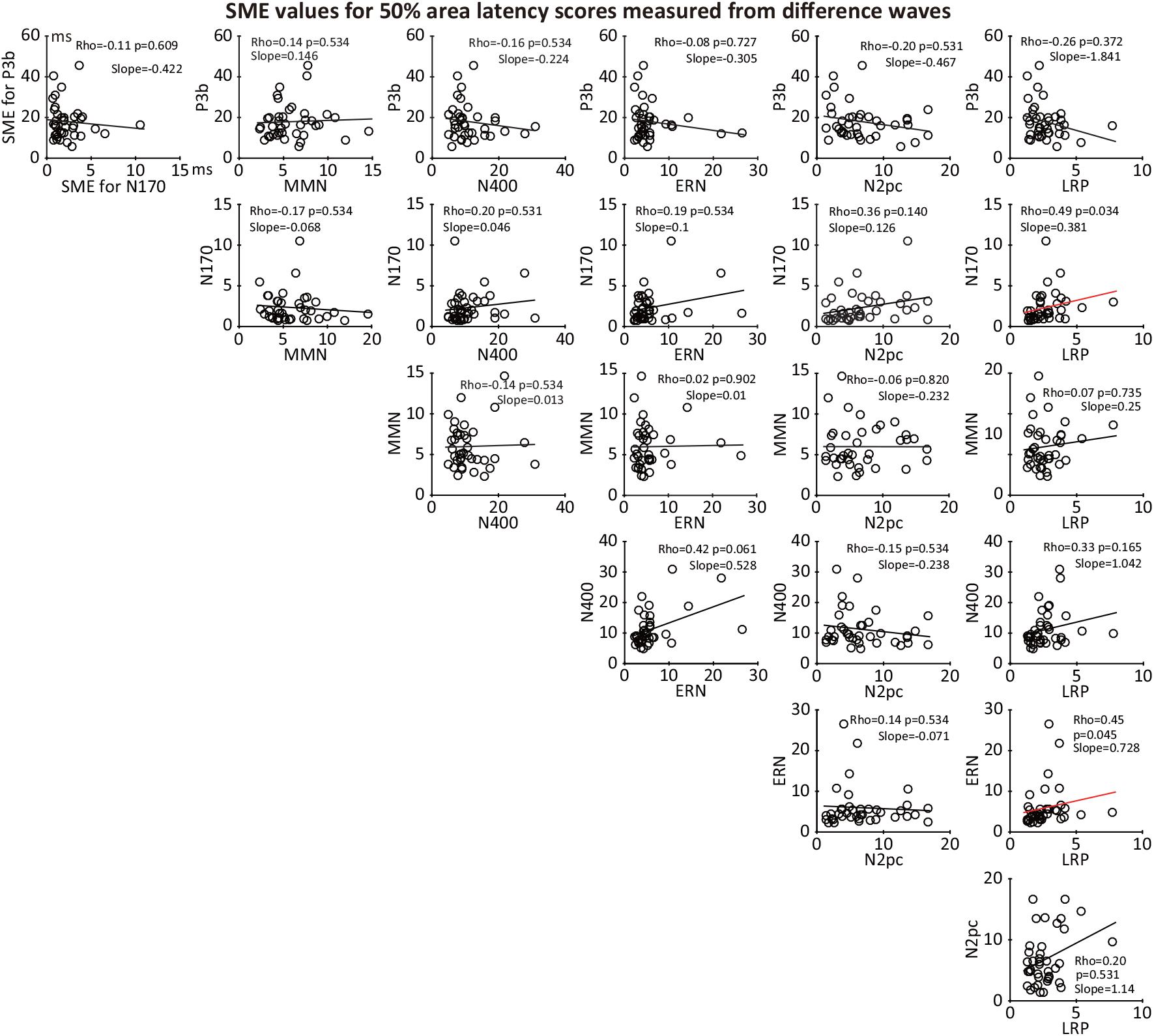
Scatterplots of the relationship between each pair of components for the SME values obtained for 50% area latency from the difference waves. Each circle represents a single participant. The p values were corrected for multiple comparisons across this entire family of tests. Note that Subject 7 was excluded from all correlations because the number of trials for one of conditions was zero for this participant in one condition of the N2pc paradigm.

**Figure S8.**
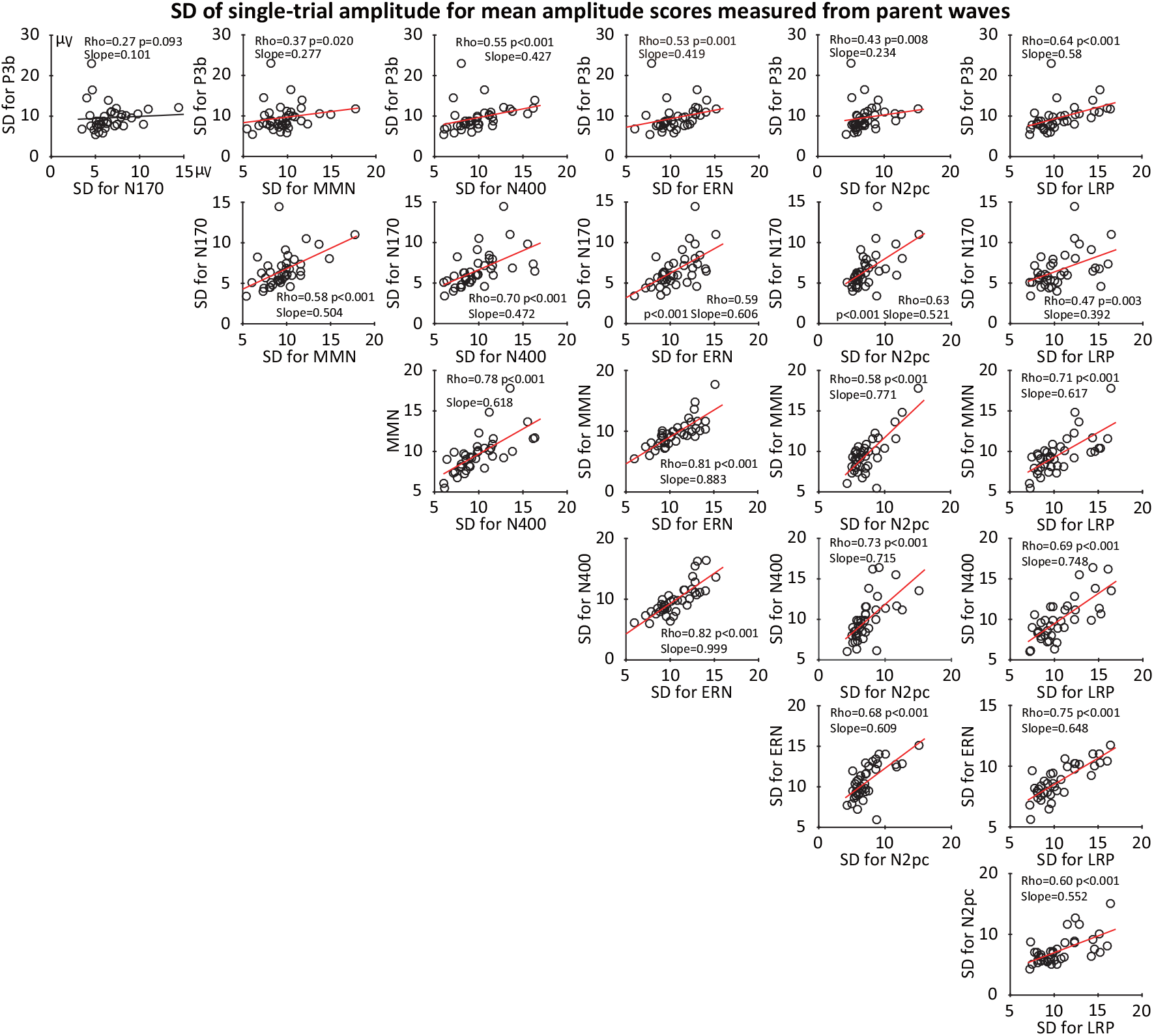
Scatterplots of the relationship between standard deviation (SD) values for mean amplitude for each pair of ERP components. The SD was calculated by measuring the mean amplitude from the single-trial epochs for a given condition and taking the SD of these values. We then averaged the SD across the two conditions used to define each component (because the SD values for the two conditions were strongly correlated, as shown in Figure S10). Each circle represents a single participant. The p values were corrected for multiple comparisons across this entire family of tests. Note that Subject 7 was excluded from all correlations because the number of trials for one of conditions was zero for this participant in one condition of the N2pc paradigm.

**Figure S9.**
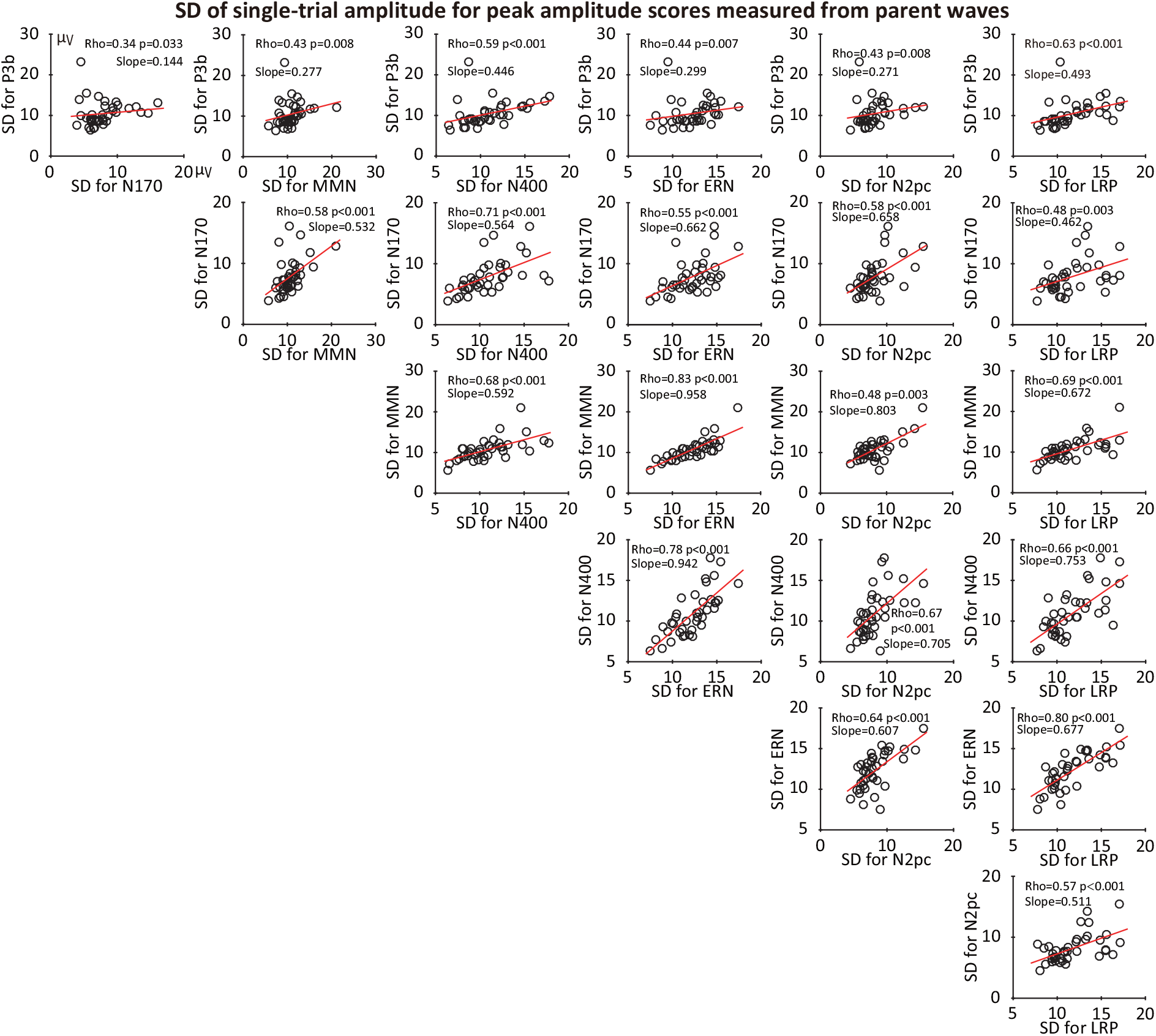
Scatterplots of the relationship between standard deviation (SD) values for peak amplitude for each pair of ERP components. The SD was calculated by measuring the mean amplitude from the single-trial epochs for a given condition and taking the SD of these values. We then averaged the SD across the two conditions used to define each component (because the SD values for the two conditions were strongly correlated, as shown in Figure S10). Each circle represents a single participant. The p values were corrected for multiple comparisons across this entire family of tests. Note that Subject 7 was excluded from all correlations because the number of trials for one of conditions was zero for this participant in one condition of the N2pc paradigm.

**Figure S10.**
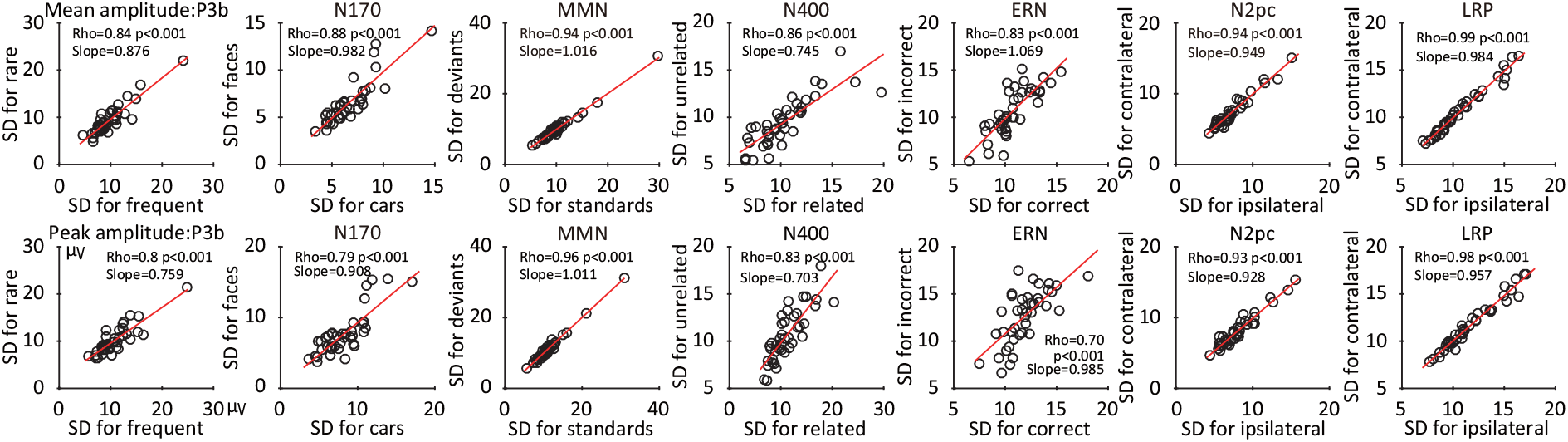
Scatterplots of the relationship between the SD values in the two conditions used to define each of the seven ERP components. The SD values were obtained from single-trial measurements of mean amplitude (top row) and peak amplitude (bottom row). Each circle represents a single participant. The p values were corrected for multiple comparisons across this entire family of tests.

## Footnotes

1 The conceptual framework described in this paper can easily be generalized to many other kinds of analyses that involve averaging and then scoring (such as when a time-frequency transform is applied prior to averaging and the data are scored as the mean power over some range of time points and frequencies).

2 Some studies compute measures of psychometric reliability, but this approach has several shortcomings, such as an inability to quantify data quality for individual participants (see (Luck et al., 2021)). A new variant of this approach can be applied to single-participant data (Clayson et al., 2021), but it applies only to mean amplitude and depends on the amount of true score variance across the participants in a given sample.

3 A measure can also be problematic if it is biased (i.e., produces a value that deviates consistently in a particular direction from the true value). Bias is a separate issue that will not be considered here but is discussed extensively in (Luck, 2014).

4 For both peak amplitude and peak latency, the local peak approach (Luck, 2014) was used, in which a peak is defined as the most extreme amplitude that is also more extreme than the average of the amplitudes at the surrounding time points. This avoids detecting a false peak when the voltage trends upward or downward at the edges of the measurement window, leading to a voltage that is more extreme than any other voltage in the window without being a peak in the waveform as a whole.

5 The RMS value is more useful than the mean across participants when the goal is to determine how the statistical power of a given experiment is influenced by the data quality. Specifically, high SME values have an outsized effect on statistical power, and this is captured by the RMS of the SME values. However, the goal of the present study is to provide a point of comparison with other studies, and for that purpose the mean is more convenient. We also provide histograms showing the entire distribution of SME scores.

## Notes

### Competing Interest Statement

The authors have declared no competing interest.

### Summary of Updates

FAIGURES

